# Sudden Death-Associated *KCNH2* Variants Have Opposing Effects on hERG1_NP_ Function

**DOI:** 10.1101/2025.10.23.684168

**Authors:** Francisco G. Sanchez-Conde, Matthew Goodrich, Olivia Stack, Louis Goldberg, Pamela Ruzycki, Abhilasha Jain, Eric N. Jimenez-Vazquez, David K. Jones

## Abstract

hERG1_NP_ is a non-conducting subdomain of the full-length hERG1 channel that is expressed in the nuclei of developing cardiomyocytes. From the nucleus, hERG1_NP_ modulates gating and expression of the full-length hERG1 channel. hERG1_NP_ variants are linked with sudden death in the young, but the impact of these variants on hERG1_NP_ activity has not been studied. To determine the effect of hERG1_NP_ variants on hERG1_NP_ activity, we measured hERG1_NP_ intracellular localization and modulation of membrane current from the full-length hERG1a channel in HEK293 cells. The wildtype hERG1_NP_ suppressed both hERG1a current (*I*_hERG_) and protein. We then screened six hERG1_NP_ variants for changes in either intracellular targeting or modulation of hERG1a current. Two variants, R885C and R1047L, disrupted hERG1_NP_ activity by altering nuclear transport or abolishing both *I*_hERG_ current suppression and nuclear targeting, respectively. Two additional variants, G1036D and Q1068R, enhanced hERG1_NP_ activity by depolarizing hERG1a’s voltage dependence of activation via accelerated channel deactivation and recovery from inactivation. Lastly, two variants had no effect: R1035W and R1069S. This work demonstrates that hERG1_NP_ variants associated with sudden death in the young can trigger both loss-of-function and gain-of-function effects on hERG1_NP_. Additional work is needed to identify specific pathogenic mechanisms of hERG1_NP_ variants.

## INTRODUCTION

*KCNH2* encodes the primary subunits of the voltage-gated potassium channel, hERG1, which conducts the rapid delayed rectifier potassium current (*I*_Kr_). In cardiac tissue, reduced *I*_Kr_ caused by loss-of-function *KCNH2* variants or off-target pharmacological block causes the cardiac disorder long QT syndrome (1,2). Patients with long QT syndrome are at increased risk of syncope, cardiac arrhythmia, and sudden cardiac death (1,2). LQTS-associated *KCNH2* variants are also linked with sudden infant death syndrome (SIDS), intrauterine fetal death, and sudden unexplained death in epilepsy (SUDEP), yet the mechanistic link between sudden death in the young and *KCNH2* variants is not fully understood (3-16).

hERG1 subunit abundance critically regulates human cardiac excitability. In human cardiomyocytes at least two hERG1 subunits combine to conduct cardiac *I*_Kr_, hERG1a and hERG1b (17-21). hERG1a and hERG1b are identical, save for their N-termini (17,18). The hERG1a N-terminal domain includes a Per-Arnt-Sim (PAS) domain that interacts with the cytoplasmic S4-S5 linker along with the cyclic nucleotide-binding homology domain (CNBHD) of the C-terminus to slow channel gating (22-26). The hERG1b N-terminal domain is unique and lacks a functional PAS domain. In a heterologous expression system, co-expressing hERG1a with hERG1b significantly accelerates hERG1 gating and increases current amplitude, compared to hERG1a homomeric channels. In human cardiomyocytes, changes in the relative abundance of hERG1a and hERG1b modulates *I*_Kr_ magnitude as well as action potential duration and susceptibility to arrhythmia (21,27-29).

The C-terminal cytoplasmic domain of hERG1 can be divided into proximal and distal halves. The proximal half is highly structured and includes a cyclic nucleotide-binding homology domain that critically regulates channel gating (22,30,31). *KCNH2* variants within the proximal half of the C-terminal domain disrupt hERG1 gating and surface expression leading to long QT syndrome (31,32). The distal half of the C-terminal domain is unstructured, save for a predicted coiled-coil domain predicted to span residues 1035-1073 (hERG1a numbering) (33-36). *KCNH2* variants within the distal half of the C-terminal domain are linked with SIDS and SUDEP, yet they minimally affect full-length channel function and the mechanism linking them with sudden death is unclear (5-16,37,38).

Previous work from our lab identified a *KCNH2*-encoded polypeptide, hERG1_NP_, that is targeted to the cell nucleus (39). Native hERG1_NP_ is enriched in the nuclei of immature cardiomyocytes and maps to the distal half of the C-terminal domain of the full-length hERG1 channel. Like nascent hERG1_NP_, the distal C-terminal domain is targeted to the cell nucleus where it reduces hERG1a current (*I*_hERG_) by roughly half in HEK293 cells stably expressing hERG1a. This work identified hERG1_NP_ as a novel modulator of hERG1 activity and a potential modifier of cardiac excitability. From the nucleus, hERG1_NP_ has the potential to broadly regulate cardiac physiology, thus hERG1_NP_ dysfunction could represent an undescribed mechanism of disease.

We hypothesized that *KCNH2* variants within the distal C-terminal domain may selectively disrupt hERG1_NP_ activity and, thus, could represent a novel mechanism of cellular dysfunction and human disease. Here we identify four hERG1_NP_ variants associated with sudden death in the young that trigger loss-of-function (R885C, R1047L) or gain-of-function (G1036D, Q1068R) effects on hERG1_NP_ activity. These data demonstrate for the first time that *KCNH2* variants associated with sudden death disrupt hERG1_NP_ activity and its regulation of the full-length hERG1 channel and may represent a novel mechanism of cellular dysfunction.

## RESULTS

### hERG1_NP_ reduces hERG1a current and protein levels

The molecular mechanisms linking *KCNH2* variants of the hERG1

C-terminal domain with sudden death are poorly understood. Here we measured the impact of several *KCNH2* variants associated with SIDS and/or SUDEP on hERG1_NP_ activity. We first validated our previous finding that hERG1_NP_ reduces *I*_hERG_ amplitude in HEK293-hERG1a cells (39). We transfected HEK293-hERG1a cells with constructs encoding either GFP or the distal C-terminal domain (hERG1_NP_) fused with mCitrine. As previously reported, hERG1_NP_ significantly reduced both steady-state and peak tail *I*_hERG_ compared to currents recorded from GFP-transfected cells (Fig. 1A-C).

**Figure 1.**
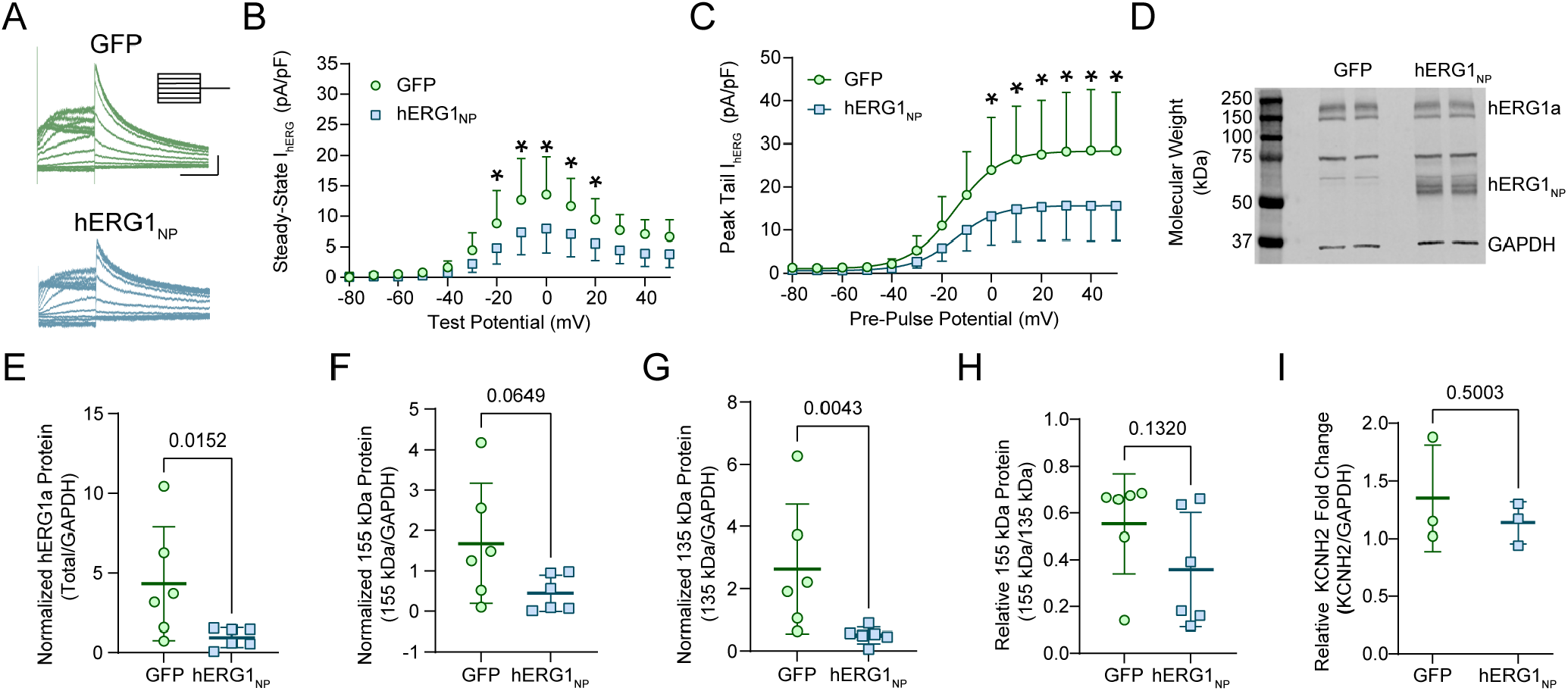
hERG1_NP_ decreases hERG1a current and protein. (A) Sample *I*_hERG_ traces elicited by the protocol in the inset from HEK293-hERG1a cells transfected with either GFP (green circles) or hERG1_NP_ (blue squares). Scale bar indicates 2 s by 5 pA/pF. (B) Steady-state hERG1a current plotted as a function of test potential from cells transfected with either GFP (green circles) or hERG1_NP_ (blue squares). (C) Peak hERG1a tail current density plotted as a function of pre-pulse potential and fitted with a Boltzmann equation (Equation 1) from HEK293-hERG1a cells transfected with either GFP (green circles) or hERG1_NP_ (blue squares). (D) Sample western blot of HEK293-hERG1a cells transfected with either GFP or hERG1_NP_ displaying bands for GAPDH, hERG1_NP_, core-glycosylated hERG1a and fully glycosylated hERG1a protein. (E) Total hERG1a protein (core-glycosylated + fully glycosylated) normalized to GAPDH from cells transfected with either GFP (green circles) or hERG1_NP_ (blue squares). (F) Fully glycosylated hERG1a protein (155 kDa) normalized to GAPDH from cells transfected with either GFP (green circles) or hERG1_NP_ (blue squares). (G) Core-glycosylated hERG1a protein (135 kDa) normalized to GAPDH from cells transfected with either GFP (green circles) or hERG1_NP_ (blue squares). (H) Quantification of 155 kDa hERG1a protein relative to 135 kDa hERG1a protein from cells transfected with either GFP (green circles) or hERG1_NP_ (blue squares). (I) Quantitative RT-PCR of *KCNH2* transcripts normalized to GAPDH from HEK293-hERG1a cells transfected with either GFP (green circles) or hERG1_NP_ (blue squares). Data in B and C were compared using a two-way ANOVA and post-hoc Bonferroni multiple comparisons test. Data in E-I were compared using a nonparametric two-tailed Mann-Whitney test. All error bars represent mean ± SD. N = 2, n = 11 for patch-clamp. N = 3, n = 6 for western blot. N = 3, n = 3 for quantitative RT-PCR. * indicates p < 0.05.

The reduced hERG1a tail current in the presence of the polypeptide suggests that hERG1_NP_ reduces total protein of the full-length hERG1a channel. To determine if hERG1_NP_ affected protein abundance, we measured hERG1a protein levels by western blot in HEK293-hERG1a cells transfected with either GFP or hERG1_NP_. Both core-glycosylated and fully glycosylated hERG1a protein bands were visible at 135 kDa and 155 kDa, respectively (Fig. 1D). Total hERG1a protein, measured as the 135 kDa signal and 155 kDa signal combined, was significantly reduced in hERG1_NP_-transfected cells compared to GFP-transfected cells (Fig. 1E). Fully glycosylated hERG1a is associated with mature hERG1 channels localized to the surface membrane, whereas core-glycosylated hERG1 represents immature hERG1 channels within the protein trafficking pathway (40-42). To determine if hERG1_NP_ affected hERG1a trafficking, we measured the abundance of the mature and immature hERG1a bands independently. In the presence of hERG1_NP_, the mature hERG1a protein band showed a downward trend that was not statistically significant (p = 0.065, Fig. 1F), whereas immature hERG1a protein levels were significantly reduced (p = 0.004, Fig. 1G). Next, we assessed the impact of hERG1_NP_ on the relative abundance of mature and immature hERG1a. We measured relative mature hERG1a as: Mature / (Mature + Immature). The relative abundance of mature protein was not significantly different compared to GFP-transfected cells (p = 0.132, Fig. 1H), suggesting that both mature and immature hERG1 are downregulated by hERG1_NP_.

To determine if the reduced hERG1a protein by hERG1_NP_ was due to a change in mRNA levels, we measured hERG1a mRNA by qRT-PCR in GFP and hERG1_NP_-transfected cells. Unlike hERG1a protein, hERG1_NP_ did not affect hERG1a transcript levels, compared to GFP controls (Fig. 1I). Collectively, the data in Figure 1 demonstrate that hERG1_NP_ reduces *I*_hERG_ and total hERG1a protein without affecting hERG1a mRNA levels.

### *KCNH2* variants alter hERG1_NP_ modulation of hERG1a current

While clinical studies have identified *KCNH2* variants in cases of SIDS and SUDEP, many variants associated with sudden death do not significantly affect full-length hERG1 channel activity (5-16,37,38). We hypothesized that variants of the distal C-terminal domain may instead selectively impact hERG1_NP_ activity. To determine if SIDS/SUDEP-associated variants of the distal C-terminal domain disrupt hERG1_NP_ activity, we transfected HEK293-hERG1a cells with hERG1_NP_-encoding constructs carrying one of the following *KCNH2* variants: R885C, R1035W, G1036D, R1047L, Q1068R, and R1069S (hERG1a numbering). Variants R885C, G1036D, R1047L, and Q1068R were identified in cases of SIDS and/or SUDEP but they each have minimal effects on the gating or expression of the full-length hERG1 channel (5-16,37,38). R1035W and R1069S were identified in clinical cohorts but are not associated with cardiac disease (7,43-48). R1035W and R1069S therefore served as negative controls. Each variant is located within one of two hERG1_NP_ structural regions of interest. R885C resides in the hERG1_NP_ nuclear localization sequence (NLS), while the remaining five sit within the predicted coiled-coil structure (Fig. 2A) (33,39).

**Figure 2.**
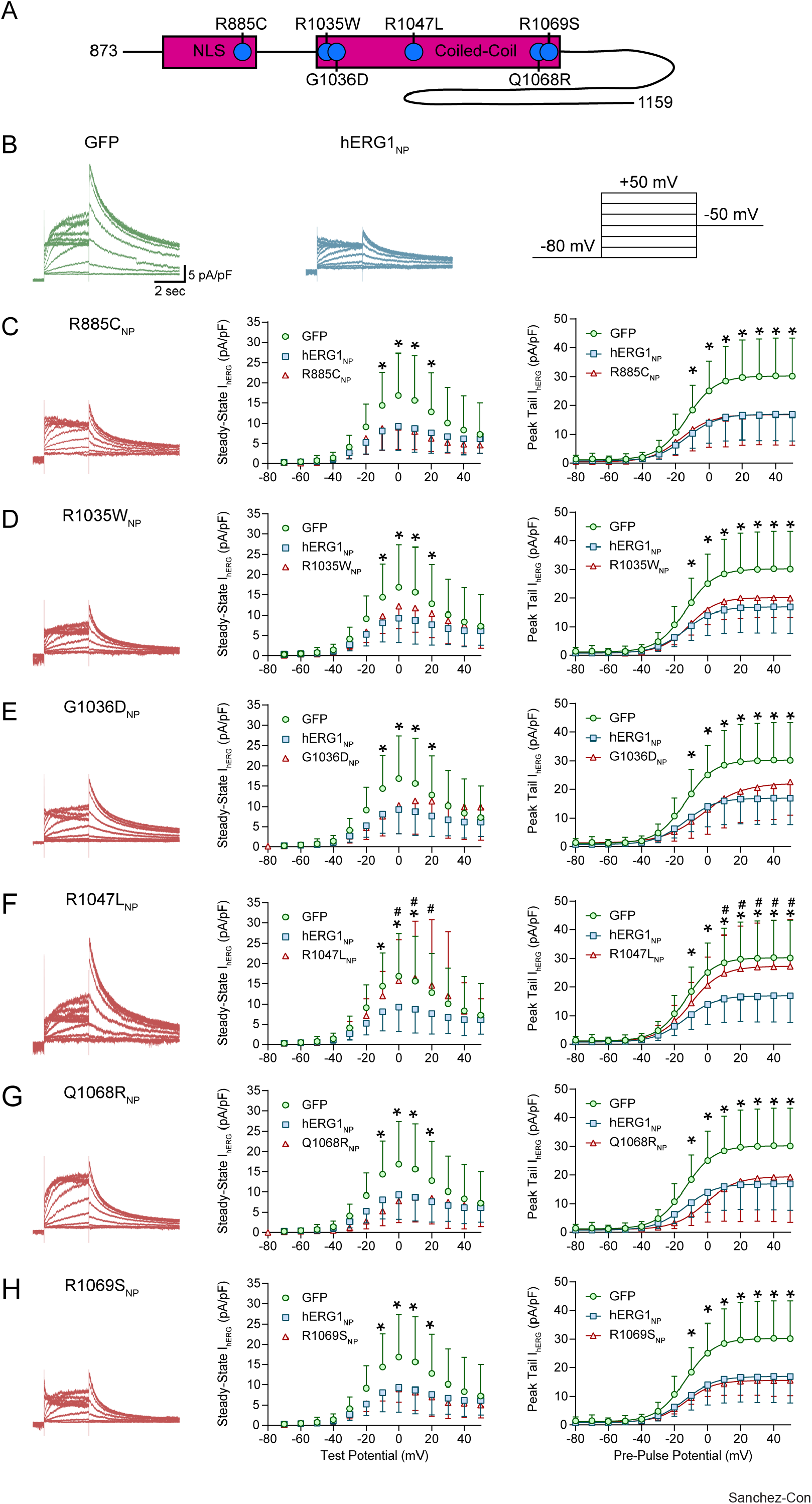
hERG1_NP_ variants differentially modulate hERG1a current. (A) Linear schematic depicting the relative location of *KCNH2* variants within hERG1_NP_. (B) Sample *I*_hERG_ traces from HEK293-hERG1a cells transfected with either GFP (green) or hERG1_NP_ (blue). (C-H) Patch clamp data from HEK293-hERG1a cells expressing GFP (green circles), hERG1_NP_ (blue squares), or hERG1_NP_ carrying one of the following *KCNH2* variants (red triangles): (C) R885C_NP_, (D) R1035W_NP_, (E) G1036D_NP_, (F) R1047L_NP_, (G) Q1068R_NP_, or (H) R1069S_NP_. Each panel displays a sample *I*_hERG_ trace recorded in the presence of the variant (left, red), steady-state current/voltage plot (middle), and peak tail current/voltage plot fitted with a Boltzmann equation (right). All variant data are compared to the same GFP and hERG1_NP_ groups but visually separated for clarity. Data in C-H were compared using a two-way ANOVA and post-hoc Dunnet multiple comparisons test with hERG1_NP_ set as the control group. Error bars represent the mean ± SD. N ≥ 3. n ≥ 7. * indicates p < 0.05 for GFP compared to hERG1_NP_. ^#^ indicates p < 0.05 for *KCNH2* variant compared to hERG1_NP_. See Table 1 for specific experimental values.

Like the wildtype polypeptide, R885C_NP_, R1035W_NP_, G1036D_NP_, Q1068R_NP_, and R1069S_NP_ all significantly reduced steady-state and peak tail current compared to recordings from GFP-transfected controls (Fig. 2B-E, G, H). In addition to reducing current amplitude, G1036D_NP_ and Q1068R_NP_ both depolarized the voltage dependence of *I*_hERG_ activation by 10 mV, compared to both wildtype hERG1_NP_ and GFP-transfected cells (Table 1; Fig. 2B, E, G). To highlight the effects of G1036D_NP_ and Q1068R_NP_ on the voltage dependence of activation, we normalized tail current amplitude to the maximum current recorded for each cell. We then plotted those data as a function of pre-pulse potential and fitted the data with a Boltzmann function (Equation 1; Fig. 3). These data demonstrate that G1036D_NP_ and Q1068R_NP_ have a gain-of-function effect on hERG1_NP_. In contrast, R1047L_NP_ had no effect on *I*_hERG_ magnitude or voltage dependence (Fig. 2B, F), suggesting that R1047L is a loss-of-function hERG1_NP_ variant. R1035W_NP_ and R1069S_NP_, which were not associated with sudden death, displayed hERG1a currents that were identical to currents recorded in the presence of wildtype hERG1_NP_. Thus, only variants associated with sudden death altered hERG1_NP_ modulation of *I*_hERG_: loss-of-function (R1047L_NP_) or gain-of-function (G1036D_NP_ and Q1068R_NP_).

**Table 1.**
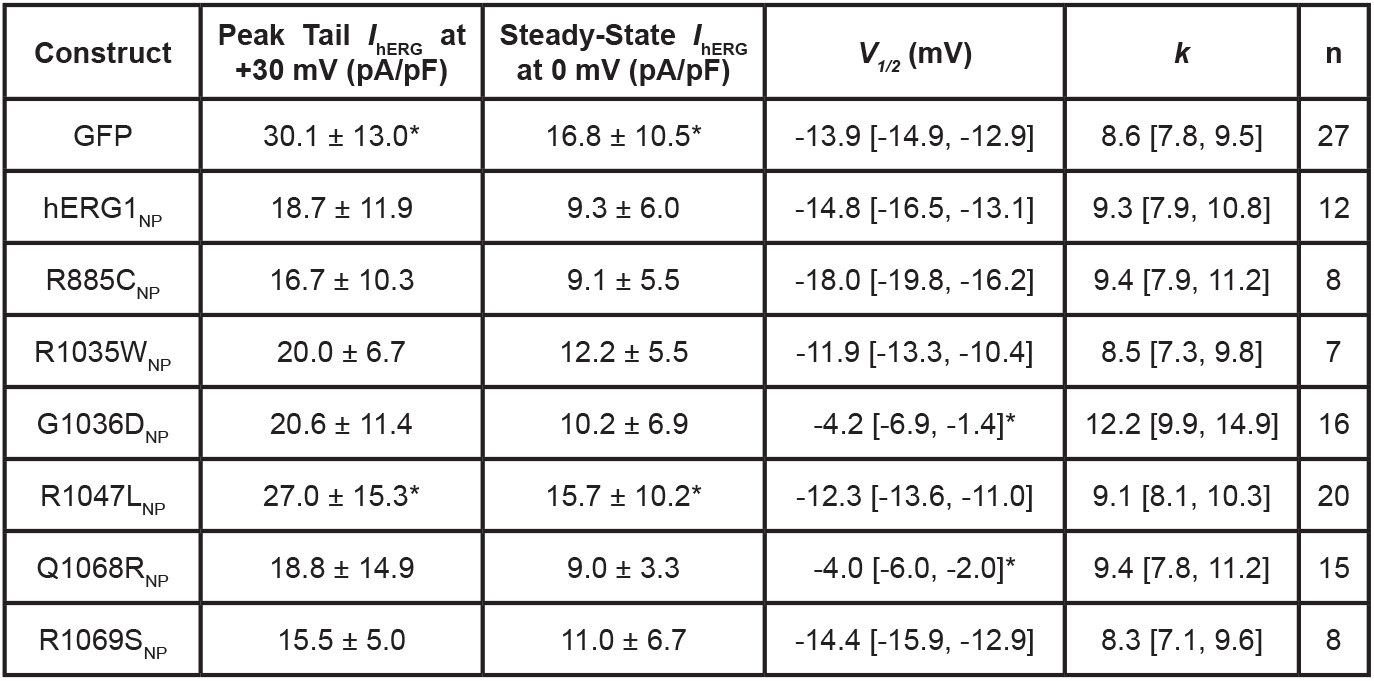
Biophysical parameters of transfected HEK293-hERG1a cells. Two-way ANOVA and post-hoc Dunnet multiple comparisons test with hERG1_NP_ set as the control group. Mean ± SD for peak tail and steady-state *I*_hERG_ magnitudes. Mean [lower limit, upper limit] for *V*_*1/2*_ and slope factor (*k*) 95% confidence intervals as calculated by GraphPad Prism. N ≥ 3. * indicates p < 0.05 compared to hERG1_NP_.

| Construct | Peak Tail $I_{\text{HERG}}$ at +30 mV (pA/pF) | Steady-State $I_{\text{HERG}}$ at 0 mV (pA/pF) | $V_{1/2}$ (mV) | $k$ | n |
| --- | --- | --- | --- | --- | --- |
| GFP | 30.1 ± 13.0* | 16.8 ± 10.5* | -13.9 [-14.9, -12.9] | 8.6 [7.8, 9.5] | 27 |
| hERG1<sub>NP</sub> | 18.7 ± 11.9 | 9.3 ± 6.0 | -14.8 [-16.5, -13.1] | 9.3 [7.9, 10.8] | 12 |
| R885C<sub>NP</sub> | 16.7 ± 10.3 | 9.1 ± 5.5 | -18.0 [-19.8, -16.2] | 9.4 [7.9, 11.2] | 8 |
| R1035W<sub>NP</sub> | 20.0 ± 6.7 | 12.2 ± 5.5 | -11.9 [-13.3, -10.4] | 8.5 [7.3, 9.8] | 7 |
| G1036D<sub>NP</sub> | 20.6 ± 11.4 | 10.2 ± 6.9 | -4.2 [-6.9, -1.4]* | 12.2 [9.9, 14.9] | 16 |
| R1047L<sub>NP</sub> | 27.0 ± 15.3* | 15.7 ± 10.2* | -12.3 [-13.6, -11.0] | 9.1 [8.1, 10.3] | 20 |
| Q1068R<sub>NP</sub> | 18.8 ± 14.9 | 9.0 ± 3.3 | -4.0 [-6.0, -2.0]* | 9.4 [7.8, 11.2] | 15 |
| R1069S<sub>NP</sub> | 15.5 ± 5.0 | 11.0 ± 6.7 | -14.4 [-15.9, -12.9] | 8.3 [7.1, 9.6] | 8 |

**Figure 3.**
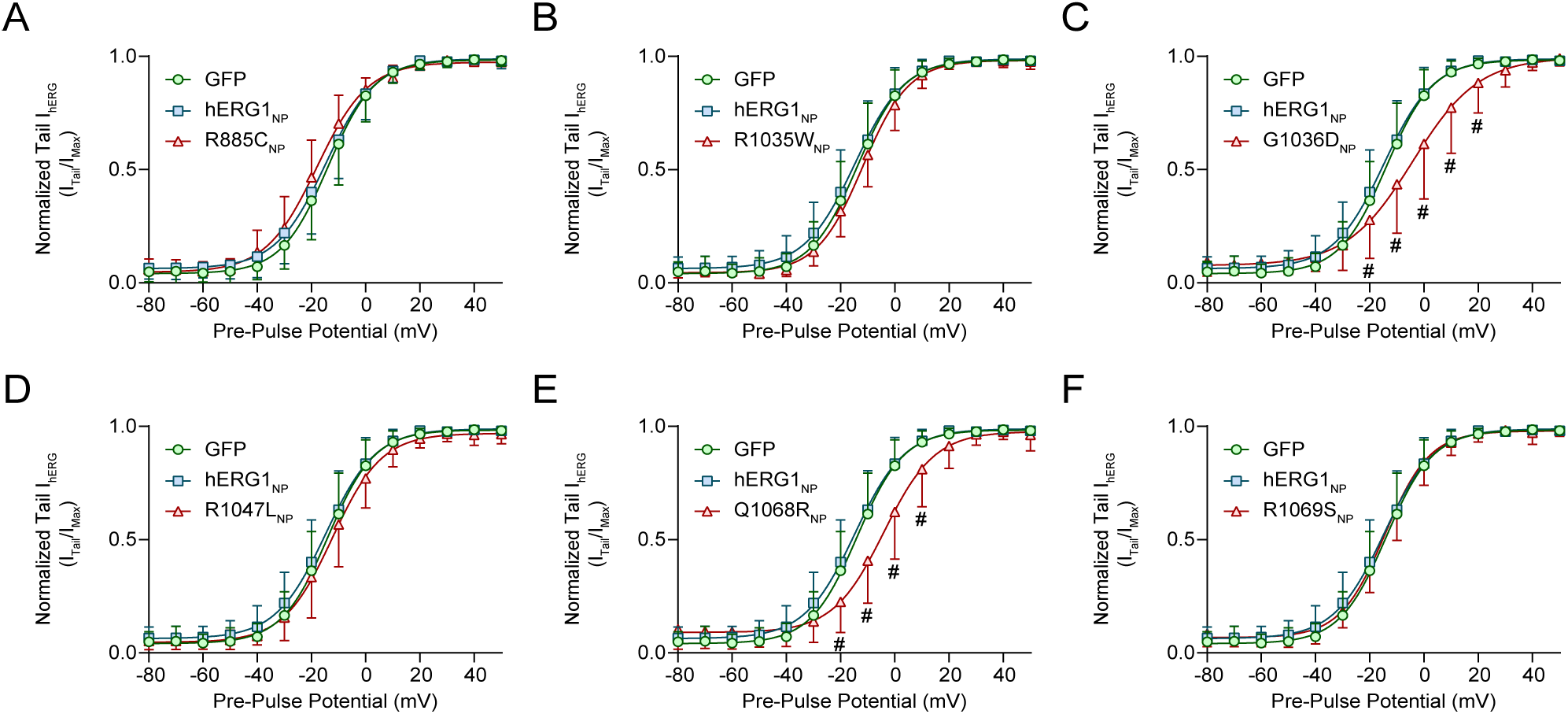
G1036D_NP_ and Q1068R_NP_ depolarize the voltage dependence of hERG1a activation. (A-F) Peak tail current/voltage relationships normalized to the maximum peak tail *I*_hERG_ for HEK293-hERG1a cells transfected with GFP (green circles), hERG1_NP_ (blue squares), or hERG1_NP_ carrying one of the six *KCNH2* variants (red triangles): (A) R885C_NP_, (B) R1035W_NP_, (C) G1036D_NP_, (D) R1047L_NP_, (E) Q1068R_NP_, and (F) R1069S_NP_. Data plotted as a function of pre-pulse potential and fitted with a Boltzmann equation (Equation 1). All variant data are compared to the same GFP and hERG1_NP_ groups but visually separated for clarity. Data in A-F were compared using a two-way ANOVA and post-hoc Dunnet multiple comparisons test with hERG1_NP_ set as the control group. Error bars represent the mean ± SD. N ≥ 3. n ≥ 7. * indicates p < 0.05 for GFP compared to hERG1_NP_. ^#^ indicates p < 0.05 for any given *KCNH2* variant compared to hERG1_NP_. See Table 1 for specific experimental values.

### G1036D_NP_ and Q1068R_NP_ accelerate the time course of deactivation

Our previous work showed that wildtype hERG1_NP_ nuclear activity slows the time course of hERG1a deactivation (39). We hypothesized that currents recorded in the presence of G1036D_NP_ and Q1068R_NP_, variants that depolarized *I*_hERG_ activation, would display accelerated deactivation compared to wildtype hERG1_NP_-transfected cells (Fig 3C, E). To test this hypothesis, we measured the time course of hERG1a deactivation by fitting current decay at -50 mV with a double exponential function (Equation 2).

Consistent with previous work, currents recorded from wildtype hERG1_NP_-transfected cells exhibited significantly larger time constants for both the fast and slow components of deactivation, compared to GFP-transfected cells (Table 2; Fig. 4A-C). G1036D_NP_ and Q1068R_NP_ both displayed accelerated rates of current decay compared to wildtype hERG1_NP_ (Table 2; Fig. 4A-C). The accelerated deactivation time constants from G1036D_NP_ and Q1068R_NP_ are consistent with the concomitant depolarization in hERG1a’s voltage dependence of activation (Fig. 3C, E). R885C_NP_, R1035W_NP_, and R1069S_NP_, which suppressed *I*_hERG_ magnitude but did not affect channel activation, displayed deactivation time constants that were comparable to wildtype hERG1_NP_ (Table 2; Fig 4B, C).

**Table 2.** Deactivation parameters of transfected HEK293-hERG1a cells. Nonparametric Kruskal-Wallis test with a post-hoc Dunn multiple comparisons correction. Comparisons included GFP vs hERG1_NP_ and all variants vs both GFP and hERG1_NP_. Mean ± SD. N ≥ 3. * indicates p < 0.05 compared to hERG1NP. ^#^ indicates p < 0.05 compared to GFP.

| Construct | Fast Tau at +30 mV (ms) | n | Slow Tau at +30 mV (ms) | n |
| --- | --- | --- | --- | --- |
| GFP | 233.8 ± 50.3* | 12 | 1727 ± 280.2* | 12 |
| hERG1<sub>NP</sub> | 393.8 ± 65.1# | 11 | 2355 ± 387.2# | 11 |
| R885C<sub>NP</sub> | 334.2 ± 45.1 | 7 | 2481 ± 412.9 | 7 |
| R1035W<sub>NP</sub> | 313.2 ± 84.8 | 7 | 1867 ± 285.4 | 7 |
| G1036D<sub>NP</sub> | 226.1 ± 51.2* | 15 | 1423 ± 261.0* | 15 |
| Q1068R<sub>NP</sub> | 237.0 ± 98.2* | 15 | 1467 ± 490.2* | 15 |
| R1069S<sub>NP</sub> | 322.1 ± 86.2 | 8 | 1914 ± 701.8 | 8 |

**Figure 4.**
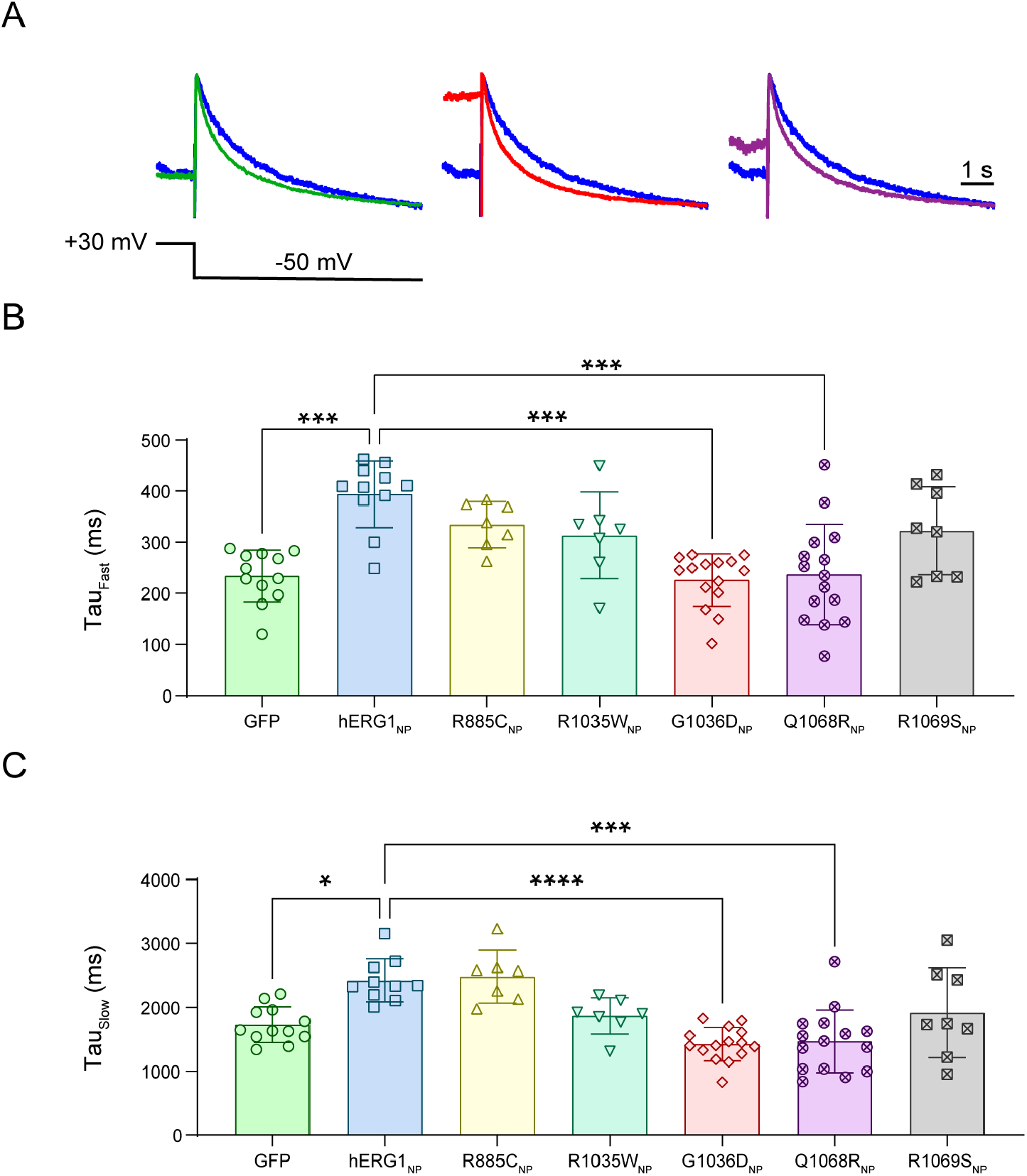
G1036D_NP_ and Q1068R_NP_ accelerate hERG1a’s time course of deactivation. (A) Sample *I*_hERG_ traces recorded from HEK293-hERG1a cells transfected with GFP (green), hERG1_NP_ (blue), G1036D_NP_ (red), or Q1068R_NP_ (purple) elicited by the displayed voltage-step protocol and scaled as indicated. Only sample traces for statistically significant variants are represented for clarity. (B) Fast and (C) slow deactivation time constants recorded from HEK293-hERG1a cells transfected with GFP (green circles), hERG1_NP_ (blue squares), R885C_NP_ (yellow upward triangles), R1035W_NP_ (aqua downward triangles), G1036D_NP_ (red diamonds), Q1068R_NP_ (purple circles) or R1069S_NP_ (grey squares). Data in B and C were compared using a nonparametric Kruskal-Wallis test with a post-hoc Dunn multiple comparisons correction. Comparisons included GFP vs hERG1_NP_ and all variants vs both GFP and hERG1_NP_. Error bars represent the mean ± SD. N ≥ 3. n ≥ 7. **** indicates p < 0.0001, *** indicates p < 0.0002, ** indicates p < 0.0021, and * indicates p < 0.0332. See Table 2 for specific experimental values.

### G1036D_NP_ and Q1068R_NP_ accelerate the time course of inactivation recovery

Having established that G1036D_NP_ and Q1068R_NP_ display altered time courses of deactivation, we tested if G1036D_NP_ and Q1068R_NP_ similarly affected hERG1a inactivation. We measured hERG1a inactivation recovery by first depolarizing the membrane with a three second conditioning pulse at 30 mV to fully activate and inactivate the channels. Cells were then stepped to a test pulse between -50 mV and -120 mV in 10 mV increments and we fitted the rebound current with a single exponential function (Equation 2). Wildtype hERG1_NP_ minimally affected inactivation recovery (Fig. 5A, B). Compared to GFP controls, hERG1_NP_ only accelerated inactivation recovery at -50 mV (Fig. 5B). In contrast, the kinetics of inactivation recovery in the presence of either G1036D_NP_ or Q1068R_NP_ were significantly faster than both GFP and wildtype hERG1_NP_, at -90 mV through -50 mV (Fig. 5A, B). The effects of G1036D_NP_ were more pronounced than Q1068R_NP_. The accelerated inactivation kinetics parallel the accelerated deactivation and depolarizing shifts in activation.

**Figure 5.**
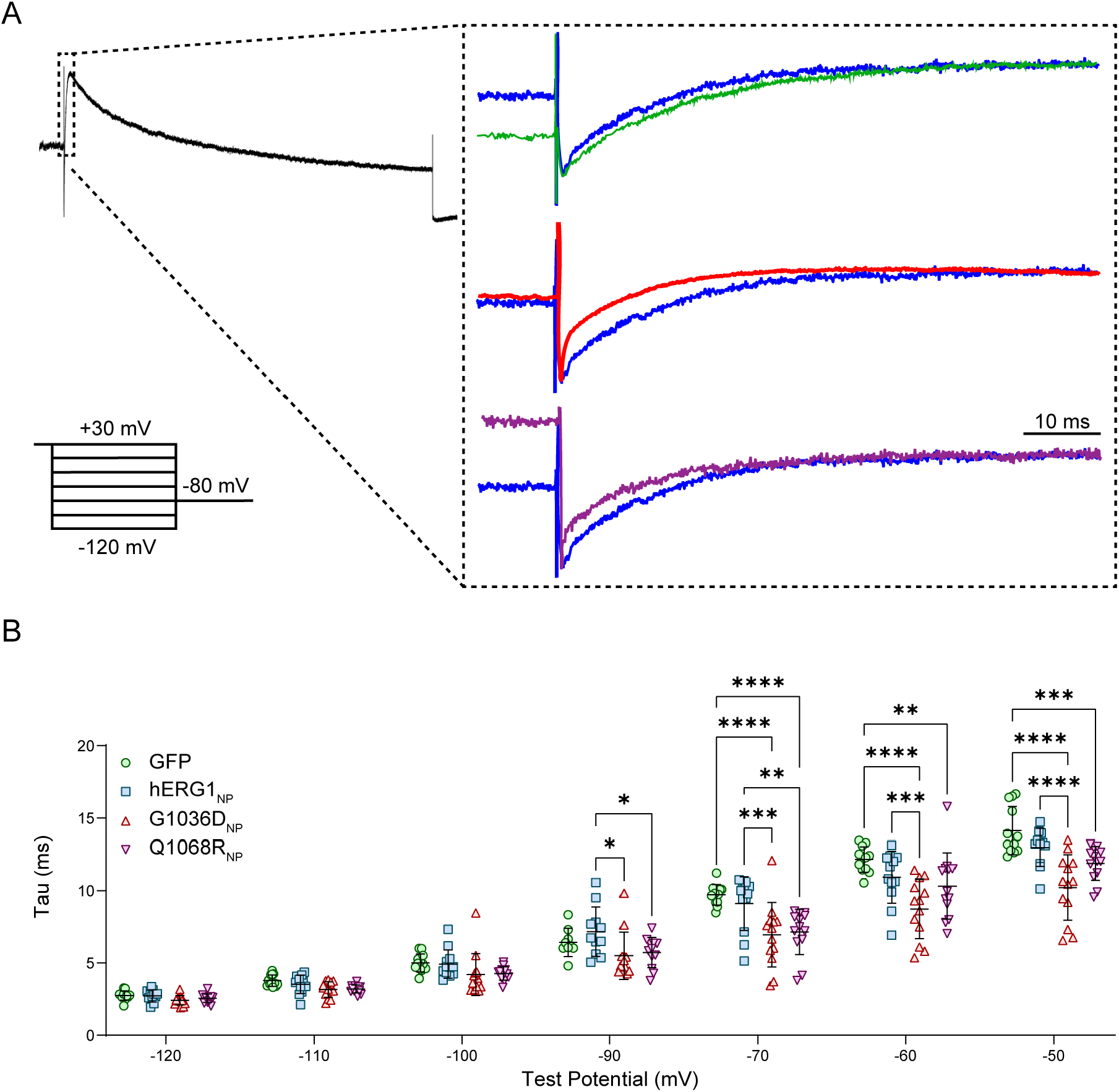
G1036D_NP_ and Q1068R_NP_ accelerate the time course of hERG1a inactivation recovery. (A) Sample *I*_hERG_ traces recorded from HEK293-hERG1a cells transfected with GFP (green), hERG1_NP_ (blue), G1036D_NP_ (red), or Q1068R_NP_ (purple) elicited by the displayed voltage-step protocol and scaled as indicated. (B) Time constants of inactivation recovery recorded from HEK293-hERG1a cells transfected with GFP (green circles), hERG1_NP_ (blue squares), G1036D_NP_ (red upward triangles), or Q1068R_NP_ (purple downward triangles). Data in B were compared using a two-way ANOVA and post-hoc Tukey multiple comparisons test. Error bars represent the mean ± SD. N = 3. n ≥ 12. **** indicates p < 0.0001, *** indicates p < 0.0002, ** indicates p < 0.0021, and * indicates p < 0.0332.

### *KCNH2* variants modulate hERG1_NP_ nuclear trafficking

Having established that R1047L_NP_, G1036D_NP_, and Q1068R_NP_ alter hERG1_NP_ modulation of *I*_hERG_, we next tested the impact of all six variants on hERG1_NP_ nuclear localization. We previously demonstrated that hERG1_NP_ modulation of hERG1a current only occurred when hERG1_NP_ was targeted to the cell’s nucleus (39). We therefore predicted that R1047L_NP_, which does not modulate *I*_hERG_, would not traffic into the nucleus. The remaining variants were predicted to retain wildtype-like nuclear localization. We transfected HEK293-hERG1a cells with GFP, hERG1_NP_, or one of the six *KCNH2* variant constructs. We also stained un-transfected HEK293-hERG1a cells with an antibody that targets the hERG1 distal C-terminal domain. The epitope of this antibody is in both the full-length hERG1 channel and hERG1_NP_ (39). We immunolabeled full-length hERG1a to demonstrate that the full-length channel alone would not produce the nuclear targeted polypeptide.

As predicted, the anti-hERG1a antibody showed robust immunofluorescence at the surface membrane and throughout the cytoplasm, but was absent from the nucleus (Fig 6A, B). In contrast, wildtype hERG1_NP_ targeted almost exclusively to the nucleus and displayed significantly greater nuclear targeting compared to GFP-transfected cells (Fig 6A, B). R1047L_NP_, which had no effect on *I*_hERG_, displayed almost exclusive cytoplasmic targeting. Surprisingly, R885C_NP_ displayed a bimodal distribution, with a roughly 50:50 split of nuclear targeted and cytoplasm targeted cells (Fig. 6A, B). R1047L_NP_’s loss of current suppression reaffirms that nuclear localization is critical for hERG1_NP_ modulation of hERG1 current. R885C_NP_’s bimodal distribution suggests that direct disruption of the hERG1_NP_ NLS causes a loss of nuclear targeting and is therefore a loss-of-function variant. The remaining constructs, R1035W_NP_, G1036D_NP_, Q1068R_NP_ and R1069S_NP_ did not modify *I*_hERG_ suppression and were targeted to the nucleus at levels comparable to wildtype hERG1_NP_ (Fig. 6A, B). These results demonstrate that a near total loss of nuclear targeting may be required to alter hERG1_NP_’s suppression of *I*_hERG_, while a partial loss of nuclear targeting is not sufficient to modify hERG1_NP_ *I*_hERG_ suppression.

**Figure 6.**
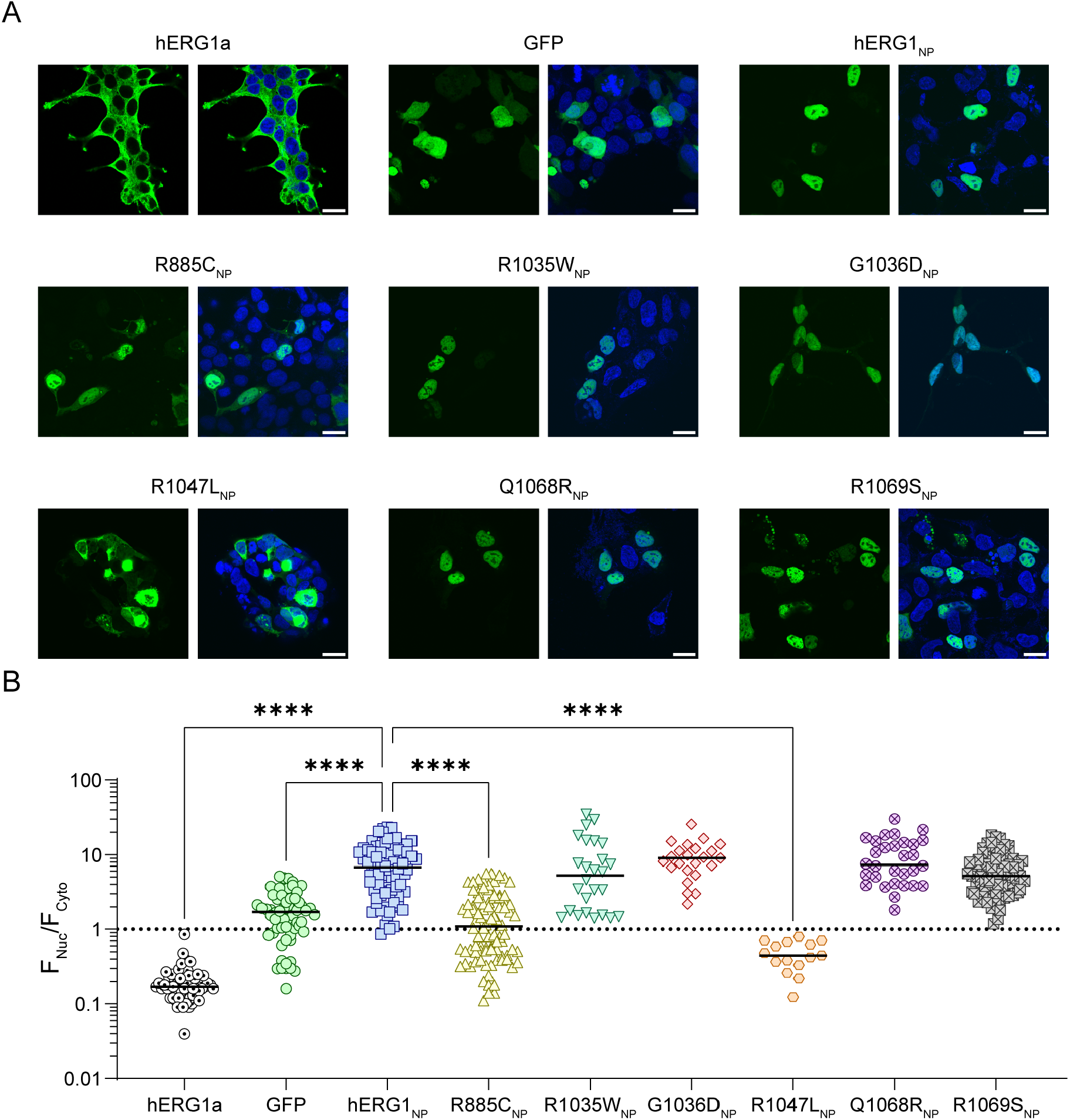
*KCNH2* variants alter hERG1_NP_ nuclear localization. (A) Sample images of un-transfected HEK293-hERG1a cells as well as HEK293-hERG1a cells transfected with GFP, hERG1_NP_, R885C_NP_, R1035W_NP_, G1036D_NP_, R1047L_NP_, Q1068R_NP_, or R1069S_NP_. Un-transfected cells are immunolabeled for the hERG1 distal C-terminal domain (green) and DAPI to delineate the nuclear (blue). Transfected groups display GFP or mCitrine fluorescence (green) and DAPI (blue). Scale bar indicates 20 µm. (B) Quantification of nuclear fluorescence intensity relative to cytoplasmic fluorescence intensity (F_Nucleus_/F_Cytoplasm_) for cells depicted in A. Data in B were compared using a nonparametric Kruskal-Wallis test with a post-hoc Dunn multiple comparisons correction and hERG1_NP_ set as the control group. Black bars represent the median. N = 3. n ≥ 15. **** indicates p < 0.0001, *** indicates p < 0.0002, ** indicates p < 0.0021, and * indicates p < 0.0332.

## DISCUSSION

The present study demonstrates that SIDS and SUDEP-associated *KCNH2* variants exert distinct effects on the nuclear-targeted hERG1_NP_ polypeptide. Consistent with our previous findings, wildtype hERG1_NP_ localizes to the nucleus and suppresses *I*_hERG_ (39), an effect associated with reduced protein levels but unchanged transcript abundance. Here, we extend these observations by showing that variants produce both gain-of-function and loss-of-function effects on hERG1_NP_ activity: G1036D_NP_ and Q1068R_NP_ enhanced gating modulation, while R885C_NP_ and R1047L_NP_ impaired nuclear targeting and/or current suppression. Together, these data provide evidence that hERG1_NP_ dysfunction may represent a novel pathogenic mechanism linking *KCNH2* variants to SIDS and SUDEP.

### *KCNH2* variant identification

The variants reported in this study were chosen to identify a potential link between hERG1_NP_ dysfunction and sudden death. R885C, which maps to the NLS of hERG1_NP_, was identified in multiple cases of SIDS but does not affect the biophysical properties of full-length hERG1 (5,6,12). The remaining variants sit within a predicted coiled-coil domain (33). R1035W was identified in several long QT patient cohorts and a non-lethal case of idiopathic ventricular fibrillation, yet R1035W has no known link with sudden death (7,43-45). G1036D was identified in several long QT study cohorts (8,43,44,49) and a case of resuscitation after cardiac arrest. Additionally, two groups found that it causes a small reduction in full-length hERG1 current, but G1036D is not predicted to be pathogenic (8,37). Like G1036D, R1047L was found in multiple LQT study cohorts (43-45); however, R1047L is also linked to cases of SIDS (5,9,11,15,16), SUDEP (10), and drug-induced torsades de pointes (38). R1047L is also present in a case of cardiac arrest resuscitation for a patient with a family history of sudden death and a separate case of posthumous sudden death in the young (SUDY) (13,14). The effects of R1047L on full-length hERG1 and its potential as a disease-causing variant are conflicting. In some hands, R1047L diminishes hERG1 current magnitude (10), in others it modifies hERG1a voltage dependence but not current magnitude (38), or the variant carries no phenotype (16,37). Bioinformatic models used for in silico analysis of R1047L are similarly inconclusive (9). Q1068R was present in multiple long QT study cohorts (43,44). It was identified in multiple cases of SIDS and functional evaluations in full-length hERG1a display small acceleration of inactivation and inactivation recovery with no change to current magnitude (11,15,16,37). Lastly, R1069S is a clinical variant with no association to sudden death (46-48). In this study we only observed changes in hERG1_NP_ activity in variants associated with sudden death, supporting the conclusion that sudden death-associated hERG1_NP_ variants alter hERG1_NP_ activity and may represent a novel mechanism of *KCNH2*-related pathophysiology.

### Gain-of-function *KCNH2* variants suggest a mechanism of hERG1a modulation

The regulatory pathways linking hERG1_NP_ nuclear activity with altered hERG1a activity remain unknown. But gain-of-function variants G1036D_NP_ and Q1068R_NP_ provide a possible insight into one mechanism by which hERG1_NP_ may regulate the full-length channel. G1036D_NP_ and Q1068R_NP_ both depolarized hERG1a’s voltage dependence of activation while accelerating the time course of deactivation and inactivation recovery. These effects were in addition to the reduced current amplitude seen by the wildtype polypeptide. These effects on hERG1a channel gating are strikingly similar to hERG1a modulation by PKA and to a lesser extent PKC.

PKA modulates hERG1 current during acute (within minutes) and chronic (>24 hours) stimulation. Multiple groups have reported that PKA phosphorylation of hERG1a initially reduces hERG1a current amplitude, accelerates deactivation, and depolarizes the voltage-dependence of activation (50,51). These effects are blocked by mutating all four of hERG1’s known PKA phosphorylation sites. Sustained PKA stimulation, using either forskolin or chlorophenyl thiol-cAMP, increases hERG1 channel synthesis leading to increased current (52,53). The effects of PKA on hERG1 voltage dependence are lost during prolonged PKA stimulation. PKC activity similarly depolarizes the voltage dependence of hERG1 activation and reduces current amplitude (54). Interestingly, PKC activity indirectly controls channel surface expression (52,53), whereas PKA directly phosphorylates hERG1 (55-57). In the case of PKC, activation leads to increased phosphorylation of Nedd4-2, the E3 ubiquitin ligase responsible for hERG1 degradation (58,59). Nedd4-2 phosphorylation inhibits ligase activity thereby reducing hERG1 degradation (50,59,60). It is possible that hERG1_NP_ nuclear activity stimulates PKA and/or PKC to regulate full-length hERG1.

### Loss-of-function *KCNH2* variants as indicators of hERG1_NP_ structure and function

R885C resides within hERG1_NP_’s NLS (39) and disrupts but does not abolish hERG1_NP_ nuclear targeting. We previously identified the karyopherin-α/β1 complex as the mediator of hERG1_NP_ trafficking (39). The results herein suggest that replacing arginine with cysteine perturbs the binding and/or recruitment of karyopherin-α, without being so disruptive as to completely nullify nuclear localization. Indeed, screening the mutated R885C_NP_ NLS with the open source software “cNLS mapper” predicts reduced but not abolished nuclear activity (61). Crystallography studies have demonstrated that positively charged lysines and arginines within any NLS form H-bonds and salt bridges with the hydrophilic residues that line the karyopherin-α binding pocket (62-68). Therefore, replacing any one of the six arginines and lysines within the hERG1_NP_ NLS would be predicted to diminish nuclear localization. R885C does not fully disrupt nuclear localization because five additional basic residues are still present within the R885C_NP_ NLS. Interestingly, despite disrupted nuclear transport, R885C_NP_ continued to suppress *I*_hERG_. This is likely due to the still substantial nuclear-localized R885C_NP_. We did not directly track the relationship between *I*_hERG_ density and magnitude of R885C_NP_ nuclear vs cytoplasmic fluorescence for individual cells. However, one might expect a bimodal distribution of *I*_hERG_ density that mimics the nuclear to cytoplasmic distribution of R885C_NP_ with high nuclear localization resulting in less current and vice versa.

R1047L resides in the hERG1_NP_ coiled-coil domain and abolished hERG1_NP_ nuclear targeting and subsequent suppression of *I*_hERG_, but the four other coiled-coil variants, R1035W, G1036D, Q1068R, and R1069S, did not affect nuclear transport. Coiled-coil domains facilitate protein-protein and protein-DNA interactions to serve an array of cellular functions (69). Many proteins form coil-mediated dimers, trimers, and tetramers to undergo nuclear trafficking making it reasonable to hypothesize that the hERG1_NP_’s coiled-coil could play a key role in maintaining the structural integrity of a hERG1_NP_ multi-polypeptide formation (70-72). R1047L replaces a polar and positively charged residue with a hydrophobic residue, which could destabilize coiled-coil assembly. However, a similarly substantial biochemical shift is observed for several additional variants that do not modify nuclear targeting: R1035W, a positive charge to a nonpolar aromatic; G1036D, a nonpolar aliphatic to a negative charge; and R1069S, a bulky positive charge to a smaller uncharged sidechain. These results may suggest that the location of the variant within the coiled-coil may also play a role in its effect on activity.

Canonical coiled-coil sequences follow a heptad repeat sequence of *abcdefg* where *a* and *d* are hydrophobic residues spaced by polar residues at *b, c, e, f*, and *g*, with *e* and *g* often carrying charges (73). As our understanding of protein structures has improved, groups have found success identifying specific residue interactions and motifs within coiled-coils that predict or trigger the higher order structural packaging of a protein (73-76). The hERG1_NP_’s predicted coiled-coil sequence follows the canonical heptad repeat motif with hydrophobic-rich *a* and *d* positions as well as charged *e* and *g* positions. For both dimeric and trimeric packaging, short coiled-coils such as the hERG1_NP_’s often carry a centrally located R-*h*-*x*-*x*-*h*-E sequence motif with an arginine residue flanked by a hydrophobic residue, two filler residues, another hydrophobic residue, and glutamic acid (74). Interestingly, this motif is selective for dimeric and trimeric but not tetrameric assemblies. However, mutation of the lead arginine to glutamine imparts tetramer-specific structural packaging, and such a Q-*h*-*x*-*x*-*h*-E motif is present within the hERG1_NP_ from residues 1048-1053 (Q-L-N-R-L-E) suggestive of tetrameric peptide packaging (77). The trimeric R-*h*-*x*-x-*h*-E motif is thought to impart structural integrity to coil interactions via key salt bridges (77). As a result, it is plausible that the hERG1_NP_’s Q-L-N-R-L-E sequence plays a similar role in tetramer stability needed for trafficking. Variant R1047L directly precedes this motif while all four other coil variants are substantially proximal or distal. Therefore, R1047L may disrupt the subsequent Q-L-N-R-L-E sequence’s interactions or spatial orientation resulting in a loss of R1047L_NP_ nuclear trafficking.

### hERG1a current as an indicator of hERG1_NP_ activity

In our initial description of hERG1_NP_ we reported that the polypeptide stabilized the hERG1a inactivated state in addition to reducing hERG1a tail current (39). In the current study, hERG1_NP_ reduced hERG1a tail current but did not affect inactivation. The discrepancy between our first study and this one highlights a limitation of using *I*_hERG_ as an indicator of hERG1_NP_. In cardiomyocytes, hERG1_NP_ nuclear targeting is upregulated in immature cells (39). This suggests that hERG1_NP_ contributes to development and stress-response pathways implicated with sudden death in the young but also demonstrates that its activity is dependent upon the physiological state of the cell in which it is expressed. Given the all too common but often unacknowledged variability of HEK293 cells, this may explain the differences between the current study and our previous work. Because hERG1_NP_ is working from within the nucleus, the downstream targets of its activity are likely dynamic and widespread. Nuclear peptides, including those derived from ion channels, are known to broadly affect transcriptional networks, chromatin remodeling, and protein assembly (78-98). We therefore acknowledge that it is likely that hERG1_NP_ mediates cellular functions undetectable through *I*_hERG_ specifically. Follow up studies identifying specific gene targets of hERG1_NP_ may help to identify more direct measures of hERG1_NP_ function and may provide a more reliable prediction of the potential pathophysiological impact of hERG1_NP_ variants.

## CONCLUSION

The experimental data presented herein demonstrate that *KCNH2* variants associated with sudden death in the young can modify hERG1_NP_ activity. While the mechanism through which hERG1_NP_ engages with the hERG1a channel remains unknown, these findings provide support for the hypothesis that hERG1_NP_ dysfunction causes human disease. hERG1_NP_ has at least one physiological output seen through hERG1a and it is reasonable to hypothesize that there is potential for hERG1_NP_ to engage with other channels or protein families across tissues or cellular compartments that require additional study.

## LIMITATIONS

Here, we report findings through experiments exclusively carried out in a line of HEK293 cells stably overexpressing the hERG1a ion channel. While our study demonstrates varied effects of *KCNH2* variants on hERG1_NP_ function, HEK293 cells are an intrinsically simplified model system that cannot replicate the complex pathways and protein dynamics that occur in specialized cell types that are known to express hERG1 protein, such as cardiomyocytes or neurons. Our system also only evaluates hERG1a homomers. In vivo, native hERG1 channels can be heterotetramers composed of at least three distinct subunits, hERG1a, hERG1b, and hERG1c (17-19,21,99-105). hERG1a contains an N-terminal Per-Arnt-Sim (PAS) domain and a C-terminal cyclic nucleotide-binding homology domain (CNBHD) (22,30,31). Channel gating is strictly regulated through direct interactions between PAS and CNBHD, and PAS and the cytoplasmic S4-S5 linker. hERG1b and hERG1c, however, carry unique N-termini that are shorter than hERG1a and lack functional PAS domains (17,18,99,102-105). Of these three subunits, hERG1a is the most well characterized and is the only one with intact PAS to CNBHD and PAS to S4-S5 linker interactions. While we chose to focus on hERG1a alone to simplify our early exploration of hERG1_NP_, the complex dynamics of different hERG1 subunits on trafficking and current introduce additional relationships to be explored in the future. Lastly, transfection of hERG1_NP_ constructs results in substantial overexpression of the polypeptides that may exacerbate the physiological effects of hERG1_NP_. Nonetheless, these data demonstrate exciting insights into the functional role and physiological relevance of hERG1_NP_ and act as an important foundation to build upon with future studies.

## MATERIALS AND METHODS

### HEK293-hERG1a cell culture

We maintained cells at 37°C with 5% CO_2_ in Heracell incubators (Thermo Fisher). We cultured HEK293 cells stably expressing hERG1a channels (HEK293-hERG1a) in Dulbecco’s Modified Eagle Medium (DMEM) supplemented with 10% fetal bovine serum (Gibco, Cat. No. 26140079) and 1% Penicillin-Streptomycin (10,000 U/mL, Gibco, Cat. No. 15140122). We split cells every 3 to 5 days at 60 to 80% confluency with PBS (Gibco, Cat. No. 10010023) and 0.05% Trypsin-EDTA (Gibco, Cat. No. 25300054). We added 50 µg/mL Geneticin antibiotic (Gibco, Cat. No. 10131035) to all freshly split cells immediately after passaging to maintain stable expression of hERG1a channels.

### DNA Constructs

The distal C-terminal domain (873-1159-mCitrine.pcDNA3) plasmid was kindly provided by Professor Matthew Trudeau at the University of Maryland Medical School. To generate the R1047L mutant, we designed forward and reverse primers with QuikChange Primer Design (agilent.com) and synthesized the primers through Eurofins (Forward: 5’-cctgttgagctggagctggagggcatc-3’, Reverse: 5’-gatgccctccagctccagctcaacagg-3’). Mutants R1035W, G1036D, Q1068R and R1069S were inserted into the distal plasmid backbone by Genscript. We verified all construct sequences with DNA sequencing provided by Eurofins.

### Quantitative RT-PCR

To measure hERG1a mRNA levels in HEK293-hERG1a cells, total RNA was prepared using the RNeasy Mini Kit (Qiagen). We synthesized cDNA by reverse transcribing 300 ng of RNA with M-MLV reverse transcriptase (Invitrogen, Cat. No. 28025-013) and oligo(dT)12–18 primers. We used IDT Mastermix (Thermo Fisher, Cat. No. 1055772) and TaqMan assay primers (10 µM, Thermo Fisher, Cat. No. 4331182 and 4351372) for GAPDH and hERG1a, respectively. We ran samples at 95°C for 30 seconds, then 39 cycles of 95°C for 3 seconds and 60°C for 20 seconds. Melting-curve analysis verified amplicon correctness. We analyzed samples in technical triplicates using a BIO-RAD C1000 Touch Thermal Cycle CFX96 (Applied Biosystems). The expression of hERG1a mRNA relative to GAPDH in cells transfected with either GFP or the distal C-terminal domain (hERG1_NP_) was calculated by the ΔΔCT method, based on the threshold cycle (CT), as fold change = 2^−(ΔΔCT), where ΔCT = CT_hERG1a_ − CT_GAPDH_ and ΔΔCT = ΔCT_hERG1NP cells_ – ΔCT_GFP cells_. From each experiment, the cDNA of 3 cell culture wells were measured as biological replicates.

### Western Blot

For protein extraction and quantification, we collected experimental cells from plastic 6-well plates by washing with ice-cold PBS (Gibco, Cat. No. 10010023) and then adding 100 µL RIPA lysis and extraction buffer (Thermo Fisher, Cat. No. 89900). We scraped cells into 1.5 mL microcentrifuge tubes and sonicated all samples before centrifuging at 12000 RPM for 15 minutes at 4°C. After centrifugation, we collected sample supernatants and diluted each 1:20 and added 5 µL per sample to a 96-well plate. We mixed DC Protein Assay Reagents A (BIO-RAD, Cat. No. 5000113) and B (BIO-RAD, Cat. No. 5000114) at a 1:50 dilution and added 200 µL of the A+B mixture to each sample well before incubating the plate at 37°C for 30 minutes. We read sample wavelengths at 490 nm to determine protein loading before diluting sample 1:1 in Laemmli sample buffer (BIO-RAD, Cat. No. 1610737). We ran 40 µg of total sample protein on a 10% SDS-PAGE gel (DI H_2_O, 30% Acrylamide, 1.5 M Tris buffer at pH 8.8, 10% SDS and TMED) topped by a stacking gel (DI H_2_O, 30% Acrylamide, 1 M Tris buffer at pH 6.8, 10% SDS and TMED). Gels ran at 80 V for 20 minutes and then 120 V for 40 minutes before transfer. We transferred protein onto nitrocellulose membranes using 200 mA in a BIO-RAD adaptor cell at 4°C. After 4 hours, we blocked with 5% nonfat dry milk in 1X TBST at room temperature for 1 hour on a countertop shaker. We then incubated nitrocellulose membranes in blocking buffer containing a 1:1000 dilution of pan-hERG C-terminal domain primary antibody (Enzo Life Sciences, Cat. No. ALX-215-049-R100) and a 1:1000 dilution of GAPDH primary antibody (Fitzgerald Industries International, Cat. No. 10RG109a) overnight at 4°C. The next day, we washed each membrane 3 times for 5 minutes per wash with 1X TBST before incubating for 1 hour at room temperature in goat anti-rabbit secondary antibody (LICORbio, Cat. No. D00115-06) and goat anti-mouse secondary antibody (LICORbio, Cat. No. 926-32210) both diluted 1:10000 in blocking buffer. We washed membranes with 1X TSBT 3 times for 5 minutes per wash before developing and quantifying blot protein signals using a LICOR Odyssey instrument.

### Immunocytochemistry

We seeded HEK293-hERG1a cells at 60% confluency on 12 mm circular glass coverslips in plastic 24-well plates. We transfected cells with 800 ng DNA 24 hours after plating using Lipofectamine 3000, P3000 (Invitrogen, Cat. No. L3000008) and Opti-MEM I Reduced Serum Medium (Gibco, Cat. No. 31985062). We replaced transfected media with fresh DMEM (10% FBS, 1% P/S) 4 to 6 hours post-transfection. After 48 hours, we fixed cells with 4% paraformaldehyde in PBS for 15 minutes. We washed the fixed cells 3 times using PBS (Gibco, Cat. No. 10010023) and stained cell nuclei using 1 μg/ml 4′,6-diamidino-2-phenylindole (DAPI, Thermo Fisher, Cat. No. 62248) for 15 minutes. For un-transfected cells, prior to DAPI staining, we immunolabeled with a 1:150 dilution of pan-hERG primary antibody (Enzo Life Sciences, Cat. No. ALX-215-049-R100) and a 1:250 dilution of goat anti-rabbit secondary antibody AF647 (Southern Biotech, Cat. No. 4050-31) to stain for the hERG1a C-terminal domain. We then washed 3 more times with PBS (Gibco, Cat. No. 10010023) and mounted the coverslips on microscope slides using ProLong™ Gold antifade reagent (Thermo Fisher, Cat. No. P36930). We completed all imaging using a Zeiss 880 confocal microscope and quantified pan-hERG antibody staining, GFP, or mCitrine localization using FIJI software.

### Electrophysiology

We seeded HEK293-hERG1a cells at ∼20% confluency as single cells in plastic 6-well plates containing 8-10 rectangular glass coverslips per well hand-cut using a glass cutter. After 24 hours, we transfected cells with 800 ng DNA using Lipofectamine 3000, P3000 (Invitrogen, Cat. No. L3000008) and Opti-MEM I Reduced Serum Medium (Gibco, Cat. No. 31985062). We replaced transfected media with fresh DMEM (10% FBS, 1% P/S) 4 to 6 hours post-transfection. We measured *I*_hERG_ using whole-cell patch-clamp at room temperature a minimum of 48 hours post-transfection.

We completed all recordings at room temperature using whole-cell patch clamp with an IPA Integrated Patch Amplifier run by SutterPatch (Sutter Instrument) within Igor Pro 8 (Wavemetrics). Data were sampled at 5 kHz and low pass filtered at 1 kHz. We recorded cells in extracellular solution containing (in mM): 150 NaCl, 5.4 KCl, 1.8 CaCl_2_, 1 MgCl_2_, 15 Glucose, 10 HEPES, 1 Na-Pyruvate and titrated to pH 7.4 using NaOH. Recording pipettes had resistances between 2 to 5 MΩ when backfilled with intracellular solution containing (in mM): 5 NaCl, 150 KCl, 2 CaCl_2_, 5 EGTA, 10 HEPES, 5 MgATP and titrated to pH 7.2 using KOH. We stored single-use 1 mL aliquots of intracellular solution at -20°C until the day of recording and kept thawed aliquots on ice throughout all recordings. All HEK293-hERG1a cells were visually confirmed to emit GFP/mCitrine fluorescence prior to single-cell recording.

To activate *I*_hERG_, we stepped cells from a holding potential of −80 mV for 1 s to a 3 s pre-pulse between −80 mV and +50 mV in 10 mV increments. Tail currents were then measured during a −50 mV, 6 s test pulse followed by a 1 s post-pulse at -80 mV. We normalized peak tail current to cellular capacitance, plotted current density as a function of pre-pulse potential in mV, and fitted the data with the following Boltzmann equation:

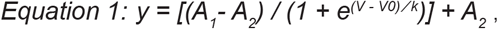

where *A*_*1*_ and *A*_*2*_ represent the maximum and minimum of the fit, respectively, *V* is the membrane potential, *V*_*0*_ is the midpoint, and *k* is the slope factor. The time course of *I*_hERG_ deactivation was assessed by fitting current decay during the test pulse with a double exponential function:

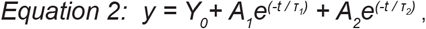

where *Y*_*0*_ is the asymptote, *A*_*1*_ and *A*_*2*_ are the relative components of the fast and slow time constants *τ*_*1*_ and *τ*_*2*_, respectively.

To evaluate the time course of inactivation recovery, we stepped cells from a holding potential of -80 mV for 1 s to a 3 s pre-pulse at +30 mV to inactivate the channels. We then measured *I*_hERG_ using 3 s test pulses stepping from -120 mV to -50 mV in 10 mV increments before stepping cells back to -80 mV for a 1 s post-pulse. We assessed *I*_hERG_ inactivation recovery by fitting an exponential function to the current immediately elicited by each respective test pulse.

### Statistical Analysis

We completed data analysis using Prism 8 (GraphPad Software, Inc) and Igor Pro 8 (Wavemetrics). We assessed values for normality (Shapiro– Wilk test) and for outlier identification (ROUT and Grubbs’ tests) before all statistical evaluation. Outliers for inactivation recovery time constants (Fig. 5) were determined as ± 2 standard deviations from the mean. Confocal imaging data are displayed as the median. Data table values are reported as mean ± SD or mean [lower limit, upper limit] 95% confidence intervals where applicable as indicated in table legends. All other data are reported as mean ± SD. Data were compared using the following statistical analysis methods, where applicable: nonparametric two-tailed Mann-Whitney test, nonparametric Kruskal-Wallis test with a Dunn multiple comparisons correction, or two-way ANOVA with Bonferroni, Dunnet, or Tukey multiple comparisons tests. We noted statistical significance at p < 0.05. Unless stated otherwise, the number “N” of observations indicates the number of transfection repeats and the number “n” of observations reflects the number of individual HEK293-hERG1a cells evaluated. We completed all experiments as single-blind studies to minimize bias.

## DATA AVAILABILITY

All data generated or analyzed in this study are included in this manuscript and its supplementary material.

## ACKNOWLEDGEMENTS

This research was supported by R01HL171039, R21NS136833, and an American SIDS Institute Grant to DKJ; T32-GM140223 to F.G.S.-C.

## REFERENCES

1. Trudeau, M. C., Warmke, J. W., Ganetzky, B., and Robertson, G. (1995) HERG, a human inward rectifier in the voltage-gated potassium channel family. Science 269, 92–95

2. Sanguinetti, M. C., Jiang, C., Curran, M. E., and Keating, M. T. (1995) A mechanistic link between an inherited and an acquired cardiac arrhythmia: HERG encodes the IKr potassium channel. Cell 81, 299–307

3. Christiansen, M., Tonder, N., Larsen, L. A., Andersen, P. S., Simonsen, H., Oyen, N., Kanters, J. K., Jacobsen, J. R., Fosdal, I., Wettrell, G., and Kjeldsen, K. (2005) Mutations in the HERG K+-ion channel: a novel link between long QT syndrome and sudden infant death syndrome. Am J Cardiol 95, 433–434

4. Crotti, L., Tester, D. J., White, W. M., Bartos, D. C., Insolia, R., Besana, A., Kunic, J. D., Will, M. L., Velasco, E. J., Bair, J. J., Ghidoni, A., Cetin, I., Van Dyke, D. L., Wick, M. J., Brost, B., Delisle, B. P., Facchinetti, F., George, A. L., Schwartz, P. J., and Ackerman, M. J. (2013) Long QT syndrome-associated mutations in intrauterine fetal death. Jama 309, 1473–1482

5. Arnestad, M., Crotti, L., Rognum, T. O., Insolia, R., Pedrazzini, M., Ferrandi, C., Vege, A., Wang, D. W., Rhodes, T. E., George, A. L., Jr., and Schwartz, P. J. (2007) Prevalence of long-QT syndrome gene variants in sudden infant death syndrome. Circulation 115, 361–367

6. Andreasen, C., Refsgaard, L., Nielsen, J. B., Sajadieh, A., Winkel, B. G., Tfelt-Hansen, J., Haunso, S., Holst, A. G., Svendsen, J. H., and Olesen, M. S. (2013) Mutations in genes encoding cardiac ion channels previously associated with sudden infant death syndrome (SIDS) are present with high frequency in new exome data. Can J Cardiol 29, 1104–1109

7. Jimenez-Jaimez, J., Alcalde Martinez, V., Jimenez Fernandez, M., Bermudez Jimenez, F., Rodriguez Vazquez Del Rey, M. D. M., Perin, F., Oyonarte Ramirez, J. M., Lopez Fernandez, S., de la Torre, I., Garcia Orta, R., Gonzalez Molina, M., Cabrerizo, E. M., Alvarez Abril, B., Alvarez, M., Macias Ruiz, R., Correa, C., and Tercedor, L. (2017) Clinical and Genetic Diagnosis of Nonischemic Sudden Cardiac Death. Rev Esp Cardiol (Engl Ed) 70, 808–816

8. Biliczki, P., Girmatsion, Z., Harenkamp, S., Anneken, L., Brandes, P., Varro, A., Marschall, C., Herrera, D., Hohnloser, S. H., Nattel, S., and Ehrlich, J. R. (2008) Cellular properties of C-terminal KCNH2 long QT syndrome mutations: description and divergence from clinical phenotypes. Heart Rhythm 5, 1159–1167

9. Glengarry, J. M., Crawford, J., Morrow, P. L., Stables, S. R., Love, D. R., and Skinner, J. R. (2014) Long QT molecular autopsy in sudden infant death syndrome. Arch Dis Child 99, 635–640

10. Soh, M. S., Bagnall, R. D., Bennett, M. F., Bleakley, L. E., Mohamed Syazwan, E. S., Marie Phillips, A., Chiam, M. D. F., McKenzie, C. E., Hildebrand, M., Crompton, D., Bahlo, M., Semsarian, C., Scheffer, E., Berkovic, S. F., and Reid, C. A. (2021) Loss-of-function variants in Kv 11.1 cardiac channels as a biomarker for SUDEP. Ann Clin Transl Neurol

11. Smith, J. L., Tester, D. J., Hall, A. R., Burgess, D. E., Hsu, C. C., Elayi, S. C., Anderson, C. L., January, C. T., Luo, J. Z., Hartzel, D. N., Mirshahi, U. L., Murray, M. F., Mirshahi, T., Ackerman, M. J., and Delisle, B. P. (2018) Functional Invalidation of Putative Sudden Infant Death Syndrome-Associated Variants in the KCNH2-Encoded Kv11.1 Channel. Circ Arrhythm Electrophysiol 11, e005859

12. Rhodes, T. E., Abraham, R. L., Welch, R. C., Vanoye, C. G., Crotti, L., Arnestad, M., Insolia, R., Pedrazzini, M., Ferrandi, C., Vege, A., Rognum, T., Roden, D. M., Schwartz, P. J., and George, A. L., Jr. (2008) Cardiac potassium channel dysfunction in sudden infant death syndrome. J Mol Cell Cardiol 44, 571–581

13. Kauferstein, S., Herz, N., Scheiper, S., Biel, S., Jenewein, T., Kunis, M., Erkapic, D., Beckmann, B. M., and Neumann, T. (2017) Relevance of molecular testing in patients with a family history of sudden death. Forensic Sci Int 276, 18–23

14. Gladding, P. A., Evans, C. A., Crawford, J., Chung, S. K., Vaughan, A., Webster, D., Neas, K., Love, D. R., Rees, M. I., Shelling, N., and Skinner, J. R. (2010) Posthumous diagnosis of long QT syndrome from neonatal screening cards. Heart Rhythm 7, 481–486

15. Tester, D. J., and Ackerman, M. J. (2005) Sudden infant death syndrome: how significant are the cardiac channelopathies? Cardiovasc Res 67, 388–396

16. Anson, B. D., Ackerman, M. J., Tester, D. J., Will, M. L., Delisle, B. P., Anderson, C. L., and January, C. T. (2004) Molecular and functional characterization of common polymorphisms in HERG (KCNH2) potassium channels. Am J Physiol Heart Circ Physiol 286, H2434–2441

17. London, B., Trudeau, M. C., Newton, K. P., Beyer, A. K., Copeland, N. G., Gilbert, D. J., Jenkins, N. A., Satler, C. A., and Robertson, G. A. (1997) Two isoforms of the mouse ether-a-go-go-related gene coassemble to form channels with properties similar to the rapidly activating component of the cardiac delayed rectifier K+ current. Circ Res 81, 870–878

18. Lees-Miller, J. P., Kondo, C., Wang, L., and Duff, H. J. (1997) Electrophysiological characterization of an alternatively processed ERG K+ channel in mouse and human hearts. Circ Res 81, 719–726

19. Jones, E. M., Roti Roti, E. C., Wang, J., Delfosse, S. A., and Robertson, G. A. (2004) Cardiac I_Kr_channels minimally comprise hERG 1a and 1b subunits. J Biol Chem 279, 44690–44694

20. Liu, F., Jones, D. K., de Lange, W. J., and Robertson, G. A. (2016) Cotranslational association of mRNA encoding subunits of heteromeric ion channels. Proc Natl Acad Sci U S A 113, 4859–4864

21. Jones, D. K., Liu, F., Vaidyanathan, R., Eckhardt, L. L., Trudeau, M. C., and Robertson, G. A. (2014) hERG 1b is critical for human cardiac repolarization. Proc Natl Acad Sci U S A 111, 18073–18077

22. Gianulis, E. C., Liu, Q., and Trudeau, M. C. (2013) Direct interaction of eag domains and cyclic nucleotide-binding homology domains regulate deactivation gating in hERG channels. J Gen Physiol 142, 351–366

23. Gustina, A. S., and Trudeau, M. C. (2013) The eag domain regulates hERG channel inactivation gating via a direct interaction. J Gen Physiol 141, 229–241

24. Gianulis, E. C., and Trudeau, M. C. (2011) Rescue of aberrant gating by a genetically encoded PAS (Per-Arnt-Sim) domain in several long QT syndrome mutant human ether-a-go-go-related gene potassium channels. J Biol Chem 286, 22160–22169

25. Trudeau, M. C., Leung, L. M., Roti, E. R., and Robertson, G. A. (2011) hERG1a N-terminal eag domain-containing polypeptides regulate homomeric hERG1b and heteromeric hERG1a/hERG1b channels: a possible mechanism for long QT syndrome. J Gen Physiol 138, 581–592

26. Gustina, A. S., and Trudeau, M. C. (2009) A recombinant N-terminal domain fully restores deactivation gating in N-truncated and long QT syndrome mutant hERG potassium channels. Proc Natl Acad Sci U S A 106, 13082–13087

27. Ukachukwu, C. U., Jimenez-Vazquez, E. N., Jain, A., and Jones, D. K. (2023) hERG1 channel subunit composition mediates proton inhibition of rapid delayed rectifier potassium current (I(Kr)) in cardiomyocytes derived from hiPSCs. J Biol Chem 299, 102778

28. Ukachukwu, C. U., Jimenez-Vazquez, E. N., Salwi, S., Goodrich, M., Sanchez-Conde, F. G., Orland, K. M., Jain, A., Eckhardt, L. L., Kamp, J., and Jones, D. K. (2025) A PAS-targeting hERG1 activator reduces arrhythmic events in Jervell and Lange-Nielsen syndrome patient-derived hiPSC-CMs. JCI Insight 10

29. Harley, C. A., Starek, G., Jones, D. K., Fernandes, A. S., Robertson, G. A., and Morais-Cabral, J. H. (2016) Enhancement of hERG channel activity by scFv antibody fragments targeted to the PAS domain. Proc Natl Acad Sci U S A 113, 9916–9921

30. Vandenberg, J. I., Torres, A. M., Campbell, T. J., and Kuchel, P. W. (2004) The HERG K+ channel: progress in understanding the molecular basis of its unusual gating kinetics. Eur Biophys J 33, 89–97

31. Codding, S. J., and Trudeau, M. C. (2019) The hERG potassium channel intrinsic ligand regulates N- and C-terminal interactions and channel closure. J Gen Physiol 151, 478–488

32. Anderson, C. L., Kuzmicki, C. E., Childs, R. R., Hintz, C. J., Delisle, B. P., and January, C. T. (2014) Large-scale mutational analysis of Kv11.1 reveals molecular insights into type 2 long QT syndrome. Nat Commun 5, 5535

33. Jenke, M., Sanchez, A., Monje, F., Stuhmer, W., Weseloh, R. M., and Pardo, L. A. (2003) C-terminal domains implicated in the functional surface expression of potassium channels. Embo J 22, 395–403

34. Wang, W., and MacKinnon, R. (2017) Cryo-EM Structure of the Open Human Ether-a-go-go-Related K(+) Channel hERG. Cell 169, 422–430 e410

35. Asai, T., Adachi, N., Moriya, T., Oki, H., Maru, T., Kawasaki, M., Suzuki, K., Chen, S., Ishii, R., Yonemori, K., Igaki, S., Yasuda, S., Ogasawara, S., Senda, T., and Murata, T. (2021) Cryo-EM Structure of K(+)-Bound hERG Channel Complexed with the Blocker Astemizole. Structure 29, 203–212 e204

36. Miyashita, Y., Moriya, T., Kato, T., Kawasaki, M., Yasuda, S., Adachi, N., Suzuki, K., Ogasawara, S., Saito, T., Senda, T., and Murata, T. (2024) Improved higher resolution cryo-EM structures reveal the binding modes of hERG channel inhibitors. Structure 32, 1926–1935 e1923

37. O’Neill, M. J., Ng, C. A., Aizawa, T., Sala, L., Bains, S., Winbo, A., Ullah, R., Shen, Q., Tan, C. Y., Kozek, K., Vanags, L. R., Mitchell, D. W., Shen, A., Wada, Y., Kashiwa, A., Crotti, L., Dagradi, F., Musu, G., Spazzolini, C., Neves, R., Bos, J. M., Giudicessi, J. R., Bledsoe, X., Gamazon, E. R., Lancaster, M. C., Glazer, A. M., Knollmann, B. C., Roden, D. M., Weile, J., Roth, F., Salem, J. E., Earle, N., Stiles, R., Agee, T., Johnson, C. N., Horie, M., Skinner, J. R., Ackerman, M. J., Schwartz, P. J., Ohno, S., Vandenberg, J. I., and Kroncke, B. M. (2024) Multiplexed Assays of Variant Effect and Automated Patch Clamping Improve KCNH2-LQTS Variant Classification and Cardiac Event Risk Stratification. Circulation 150, 1869–1881

38. Sun, Z., Milos, P. M., Thompson, J. F., Lloyd, D. B., Mank-Seymour, A., Richmond, J., Cordes, J. S., and Zhou, J. (2004) Role of a KCNH2 polymorphism (R1047 L) in dofetilide-induced Torsades de Pointes. J Mol Cell Cardiol 37, 1031–1039

39. Jain, A., Stack, O., Ghodrati, S., Sanchez-Conde, F. G., Ukachukwu, C. U., Salwi, S., Jimenez-Vazquez, E. N., and Jones, D. K. (2023) KCNH2 encodes a nuclear-targeted polypeptide that mediates hERG1 channel gating and expression. Proc Natl Acad Sci U S A 120, e2214700120

40. Zhou, Z., Gong, Q., Ye, B., Fan, Z., Makielski, J. C., Robertson, G. A., and January, C. T. (1998) Properties of HERG channels stably expressed in HEK 293 cells studied at physiological temperature. Biophys J 74, 230–241

41. Petrecca, K., Atanasiu, R., Akhavan, A., and Shrier, A. (1999) N-linked glycosylation sites determine HERG channel surface membrane expression. J Physiol 515 (Pt 1), 41–48

42. Zhang, Y., Hartmann, H. A., and Satin, J. (1999) Glycosylation influences voltage-dependent gating of cardiac and skeletal muscle sodium channels. J Membr Biol 171, 195–207

43. Kapa, S., Tester, D. J., Salisbury, B. A., Harris-Kerr, C., Pungliya, M. S., Alders, M., Wilde, A. A., and Ackerman, M. J. (2009) Genetic testing for long-QT syndrome: distinguishing pathogenic mutations from benign variants. Circulation 120, 1752–1760

44. Giudicessi, J. R., Kapplinger, J. D., Tester, D. J., Alders, M., Salisbury, B. A., Wilde, A. A., and Ackerman, M. J. (2012) Phylogenetic and physicochemical analyses enhance the classification of rare nonsynonymous single nucleotide variants in type 1 and 2 long-QT syndrome. Circ Cardiovasc Genet 5, 519–528

45. Ackerman, M. J., Tester, D. J., Jones, G. S., Will, M. L., Burrow, C. R., and Curran, M. E. (2003) Ethnic differences in cardiac potassium channel variants: implications for genetic susceptibility to sudden cardiac death and genetic testing for congenital long QT syndrome. Mayo Clin Proc 78, 1479–1487

46. Walsh, R., Thomson, K. L., Ware, J. S., Funke, B. H., Woodley, J., McGuire, K. J., Mazzarotto, F., Blair, E., Seller, A., Taylor, J. C., Minikel, E. V., Exome Aggregation, C., MacArthur, D. G., Farrall, M., Cook, S. A., and Watkins, H. (2017) Reassessment of Mendelian gene pathogenicity using 7,855 cardiomyopathy cases and 60,706 reference samples. Genet Med 19, 192–203

47. Pugh, T. J., Kelly, M. A., Gowrisankar, S., Hynes, E., Seidman, M. A., Baxter, S. M., Bowser, M., Harrison, B., Aaron, D., Mahanta, L. M., Lakdawala, N. K., McDermott, G., White, E. T., Rehm, H. L., Lebo, M., and Funke, B. H. (2014) The landscape of genetic variation in dilated cardiomyopathy as surveyed by clinical DNA sequencing. Genet Med 16, 601–608

48. Alfares, A. A., Kelly, M. A., McDermott, G., Funke, B. H., Lebo, M. S., Baxter, S. B., Shen, J., McLaughlin, H. M., Clark, E. H., Babb, L. J., Cox, S. W., DePalma, S. R., Ho, C. Y., Seidman, J. G., Seidman, C. E., and Rehm, H. L. (2015) Results of clinical genetic testing of 2,912 probands with hypertrophic cardiomyopathy: expanded panels offer limited additional sensitivity. Genet Med 17, 880–888

49. Tester, D. J., Will, M. L., Haglund, C. M., and Ackerman, M. J. (2005) Compendium of cardiac channel mutations in 541 consecutive unrelated patients referred for long QT syndrome genetic testing. Heart Rhythm 2, 507–517

50. Thomas, D., Wu, K., Wimmer, A. B., Zitron, E., Hammerling, B. C., Kathofer, S., Lueck, S., Bloehs, R., Kreye, V. A., Kiehn, J., Katus, H. A., Schoels, W., and Karle, C. A. (2004) Activation of cardiac human ether-a-go-go related gene potassium currents is regulated by alpha(1A)-adrenoceptors. J Mol Med (Berl) 82, 826–837

51. Krishnan, Y., Li, Y., Zheng, R., Kanda, V., and McDonald, T. V. (2012) Mechanisms underlying the protein-kinase mediated regulation of the HERG potassium channel synthesis. Biochim Biophys Acta 1823, 1273–1284

52. Cui, J., Melman, Y., Palma, E., Fishman, G. I., and McDonald, T. V. (2000) Cyclic AMP regulates the HERG K(+) channel by dual pathways. Curr Biol 10, 671–674

53. Wei, Z., Thomas, D., Karle, C. A., Kathofer, S., Schenkel, J., Kreye, V. A., Ficker, E., Wible, B. A., and Kiehn, J. (2002) Protein kinase A-mediated phosphorylation of HERG potassium channels in a human cell line. Chin Med J (Engl) 115, 668–676

54. Shu, L., Zhang, W., Su, G., Zhang, J., Liu, C., and Xu, J. (2013) Modulation of HERG K+ channels by chronic exposure to activators and inhibitors of PKA and PKC: actions independent of PKA and PKC phosphorylation. Cell Physiol Biochem 32, 1830–1844

55. Chen, J., Sroubek, J., Krishnan, Y., Li, Y., Bian, J., and McDonald, T. V. (2009) PKA phosphorylation of HERG protein regulates the rate of channel synthesis. Am J Physiol Heart Circ Physiol 296, H1244–1254

56. Chen, J., Chen, K., Sroubek, J., Wu, Z. Y., Thomas, D., Bian, J. S., and McDonald, T. V. (2010) Post-transcriptional control of human ether-a-go-go-related gene potassium channel protein by alpha-adrenergic receptor stimulation. Mol Pharmacol 78, 186–197

57. Sroubek, J., and McDonald, T. V. (2011) Protein kinase A activity at the endoplasmic reticulum surface is responsible for augmentation of human ether-a-go-go-related gene product (HERG). J Biol Chem 286, 21927–21936

58. Albesa, M., Grilo, L. S., Gavillet, B., and Abriel, H. (2011) Nedd4-2-dependent ubiquitylation and regulation of the cardiac potassium channel hERG1. J Mol Cell Cardiol 51, 90–98

59. Sutherland-Deveen, M. E., Wang, T., Lamothe, S. M., Tschirhart, J. N., Guo, J., Li, W., Yang, T., Du, Y., and Zhang, S. (2019) Differential Regulation of Human Ether-a-Go-Go-Related Gene (hERG) Current and Expression by Activation of Protein Kinase C. Mol Pharmacol 96, 1–12

60. Cockerill, S. L., Tobin, A. B., Torrecilla, I., Willars, G. B., Standen, N. B., and Mitcheson, J. S. (2007) Modulation of hERG potassium currents in HEK-293 cells by protein kinase C. Evidence for direct phosphorylation of pore forming subunits. J Physiol

61. Kosugi, S., Hasebe, M., Tomita, M., and Yanagawa, H. (2009) Systematic identification of cell cycle-dependent yeast nucleocytoplasmic shuttling proteins by prediction of composite motifs. Proc Natl Acad Sci U S A 106, 10171–10176

62. Lange, A., Mills, R. E., Lange, C. J., Stewart, M., Devine, S. E., and Corbett, A. H. (2007) Classical nuclear localization signals: definition, function, and interaction with importin alpha. J Biol Chem 282, 5101–5105

63. Dingwall, C., and Laskey, R. A. (1991) Nuclear targeting sequences--a consensus? Trends Biochem Sci 16, 478–481

64. Robbins, J., Dilworth, S. M., Laskey, R. A., and Dingwall, C. (1991) Two interdependent basic domains in nucleoplasmin nuclear targeting sequence: identification of a class of bipartite nuclear targeting sequence. Cell 64, 615–623

65. Kalderon, D., Richardson, W. D., Markham, A. F., and Smith, A. E. (1984) Sequence requirements for nuclear location of simian virus 40 large-T antigen. Nature 311, 33–38

66. Conti, E., and Kuriyan, J. (2000) Crystallographic analysis of the specific yet versatile recognition of distinct nuclear localization signals by karyopherin alpha. Structure 8, 329–338

67. Fontes, M. R., Teh, T., and Kobe, B. (2000) Structural basis of recognition of monopartite and bipartite nuclear localization sequences by mammalian importin-alpha. J Mol Biol 297, 1183–1194

68. Hodel, M. R., Corbett, A. H., and Hodel, A. E. (2001) Dissection of a nuclear localization signal. J Biol Chem 276, 1317–1325

69. Truebestein, L., and Leonard, T. A. (2016) Coiled-coils: The long and short of it. Bioessays 38, 903–916

70. Ludwig, J., Owen, D., and Pongs, O. (1997) Carboxy-terminal domain mediates assembly of the voltage-gated rat ether-à-go-go potassium channel. Embo J 16, 6337–6345

71. Kupershmidt, S., Snyders, D. J., Raes, A., and Roden, D. M. (1998) A K+ channel splice variant common in human heart lacks a C-terminal domain required for expression of rapidly activating delayed rectifier current. J Biol Chem 273, 27231–27235

72. Kupershmidt, S., Yang, T., Chanthaphaychith, S., Wang, Z., Towbin, J. A., and Roden, D. M. (2002) Defective human Ether-a-go-go-related gene trafficking linked to an endoplasmic reticulum retention signal in the C terminus. J Biol Chem 277, 27442–27448

73. Woolfson, D. N. (2023) Understanding a protein fold: The physics, chemistry, and biology of alpha-helical coiled coils. J Biol Chem 299, 104579

74. Ciani, B., Bjelic, S., Honnappa, S., Jawhari, H., Jaussi, R., Payapilly, A., Jowitt, T., Steinmetz, M. O., and Kammerer, R. A. (2010) Molecular basis of coiled-coil oligomerization-state specificity. Proc Natl Acad Sci U S A 107, 19850–19855

75. Steinmetz, M. O., Stock, A., Schulthess, T., Landwehr, R., Lustig, A., Faix, J., Gerisch, G., Aebi, U., and Kammerer, R. A. (1998) A distinct 14 residue site triggers coiled-coil formation in cortexillin I. EMBO J 17, 1883–1891

76. Steinmetz, M. O., Jelesarov, I., Matousek, W. M., Honnappa, S., Jahnke, W., Missimer, J. H., Frank, S., Alexandrescu, A. T., and Kammerer, R. A. (2007) Molecular basis of coiled-coil formation. Proc Natl Acad Sci U S A 104, 7062–7067

77. Beck, K., Gambee, J. E., Kamawal, A., and Bachinger, H. P. (1997) A single amino acid can switch the oligomerization state of the alpha-helical coiled-coil domain of cartilage matrix protein. EMBO J 16, 3767–3777

78. Kim, K. H., Son, J. M., Benayoun, B. A., and Lee, C. (2018) The Mitochondrial-Encoded Peptide MOTS-c Translocates to the Nucleus to Regulate Nuclear Gene Expression in Response to Metabolic Stress. Cell Metab 28, 516–524 e517

79. Mangalhara, K. C., and Shadel, G. S. (2018) A Mitochondrial-Derived Peptide Exercises the Nuclear Option. Cell Metab 28, 330–331

80. Yang, Y., Gao, H., Zhou, H., Liu, Q., Qi, Z., Zhang, Y., and Zhang, J. (2019) The role of mitochondria-derived peptides in cardiovascular disease: Recent updates. Biomed Pharmacother 117, 109075

81. Reits, E., Griekspoor, A., Neijssen, J., Groothuis, T., Jalink, K., van Veelen, P., Janssen, H., Calafat, J., Drijfhout, J. W., and Neefjes, J. (2003) Peptide diffusion, protection, and degradation in nuclear and cytoplasmic compartments before antigen presentation by MHC class I. Immunity 18, 97–108

82. Fedoreyeva, L. I., Vanyushin, B. F., and Baranova, E. N. (2020) Peptide AEDL alters chromatin conformation via histone binding. Aims Biophys 7, 1–16

83. Teles, K., Fernandes, V., Silva, I., Leite, M., Grisolia, C., Lobbia, V. R., van Ingen, H., Honorato, R., Lopes-de-Oliveira, P., Treptow, W., and Santos, G. (2020) Nucleosome binding peptide presents laudable biophysical and in vivo effects. Biomed Pharmacother 121, 109678

84. Janssens, Y., Wynendaele, E., Vanden Berghe, W., and De Spiegeleer, B. (2019) Peptides as epigenetic modulators: therapeutic implications. Clin Epigenetics 11, 101

85. Yang, Y., Yu, Z., Geng, J., Liu, M., Liu, N., Li, P., Hong, W., Yue, S., Jiang, H., Ge, H., Qian, F., Xiong, W., Wang, P., Song, S., Li, X., Fan, Y., and Liu, X. (2022) Cytosolic peptides encoding Ca(V)1 C-termini downregulate the calcium channel activity-neuritogenesis coupling. Commun Biol 5, 484

86. Gomez-Ospina, N., Tsuruta, F., Barreto-Chang, O., Hu, L., and Dolmetsch, R. (2006) The C terminus of the L-type voltage-gated calcium channel Ca(V)1.2 encodes a transcription factor. Cell 127, 591–606

87. Liu, X., Yang, P. S., Yang, W., and Yue, D. T. (2010) Enzymeinhibitor-like tuning of Ca(2+) channel connectivity with calmodulin. Nature 463, 968–972

88. Singh, A., Hamedinger, D., Hoda, J. C., Gebhart, M., Koschak, A., Romanin, C., and Striessnig, J. (2006) C-terminal modulator controls Ca2+-dependent gating of Ca(v)1.4 L-type Ca2+ channels. Nat Neurosci 9, 1108–1116

89. Hulme, J. T., Yarov-Yarovoy, V., Lin, T. W., Scheuer, T., and Catterall, W. A. (2006) Autoinhibitory control of the CaV1.2 channel by its proteolytically processed distal C-terminal domain. J Physiol 576, 87–102

90. Morrill, J. A., and Cannon, S. C. (2000) COOH-terminal truncated alpha(1S) subunits conduct current better than full-length dihydropyridine receptors. J Gen Physiol 116, 341–348

91. Sang, L., Dick, I. E., and Yue, D. T. (2016) Protein kinase A modulation of CaV1.4 calcium channels. Nat Commun 7, 12239

92. Bock, G., Gebhart, M., Scharinger, A., Jangsangthong, W., Busquet, P., Poggiani, C., Sartori, S., Mangoni, M. E., Sinnegger-Brauns, M. J., Herzig, S., Striessnig, J., and Koschak, A. (2011) Functional properties of a newly identified C-terminal splice variant of Cav1.3 L-type Ca2+ channels. J Biol Chem 286, 42736–42748

93. Wei, X., Neely, A., Lacerda, A. E., Olcese, R., Stefani, E., Perez-Reyes, E., and Birnbaumer, L. (1994) Modification of Ca2+ channel activity by deletions at the carboxyl terminus of the cardiac alpha 1 subunit. J Biol Chem 269, 1635–1640

94. Gerhardstein, B. L., Gao, T., Bunemann, M., Puri, T. S., Adair, A., Ma, H., and Hosey, M. M. (2000) Proteolytic processing of the C terminus of the alpha(1C) subunit of L-type calcium channels and the role of a proline-rich domain in membrane tethering of proteolytic fragments. J Biol Chem 275, 8556–8563

95. Sang, L., Vieira, D. C. O., Yue, D. T., Ben-Johny, M., and Dick, I. E. (2021) The molecular basis of the inhibition of Ca(V)1 calciumdependent inactivation by the distal carboxy tail. J Biol Chem 296, 100502

96. Schroder, E., Byse, M., and Satin, J. (2009) L-type calcium channel C terminus autoregulates transcription. Circ Res 104, 1373–1381

97. Lu, L., Sirish, P., Zhang, Z., Woltz, R. L., Li, N., Timofeyev, V., Knowlton, A. A., Zhang, X. D., Yamoah, E. N., and Chiamvimonvat, N. (2015) Regulation of gene transcription by voltage-gated L-type calcium channel, Cav1.3. J Biol Chem 290, 4663–4676

98. Krapivinsky, G., Krapivinsky, L., Manasian, Y., and Clapham, D. E. (2014) The TRPM7 chanzyme is cleaved to release a chromatin-modifying kinase. Cell 157, 1061–1072

99. London, B., Aydar, E., Lewarchik, C., Seibel, J. S., January, C. T., and Robertson, G. A. (1998) N and C-terminal isoforms of HERG in the human heart. Biophysical J.

100. Splawski, I., Shen, J., Timothy, K. W., Vincent, G. M., Lehmann, M. H., an Keating, M. T. (1998) Genomic structure of three long QT syndrome genes: KVLQT1, HERG, and KCNE1. Genomics 51, 86–97

101. Sale, H., Wang, J., O’Hara, T. J., Tester, D. J., Phartiyal, P., He, J. Q., Rudy, Y., Ackerman, M. J., and Robertson, G. A. (2008) Physiological properties of hERG 1a/1b heteromeric currents and a hERG 1b-specific mutation associated with Long-QT syndrome. Circ Res 103, e81–95

102. Huffaker, S. J., Chen, J., Nicodemus, K. K., Sambataro, F., Yang, F., Mattay, V., Lipska, B. K., Hyde, T. M., Song, J., Rujescu, D., Giegling, I., Mayilyan, K., Proust, M. J., Soghoyan, A., Caforio, G., Callicott, J. H., Bertolino, A., Meyer-Lindenberg, A., Chang, J., Ji, Y., Egan, M. F., Goldberg, T. E., Kleinman, J. E., Lu, B., and Weinberger, D. R. (2009) A primate-specific, brain isoform of KCNH2 affects cortical physiology, cognition, neuronal repolarization and risk of schizophrenia. Nat Med 15, 509–518

103. Calcaterra, N. E., Hoeppner, D. J., Wei, H., Jaffe, A. E., Maher, B. J., and Barrow, J. C. (2016) Schizophrenia-Associated hERG channel Kv11.1-3.1 Exhibits a Unique Trafficking Deficit that is Rescued Through Proteasome Inhibition for High Throughput Screening. Sci Rep 6, 19976

104. Carr, G. V., Chen, J., Yang, F., Ren, M., Yuan, P., Tian, Q., Bebensee, A., Zhang, G. Y., Du, J., Glineburg, P., Xun, R., Akhile, O., Akuma, D., Pickel, J., Barrow, J. C., Papaleo, F., and Weinberger, D. R. (2016) KCNH2-3.1 expression impairs cognition and alters neuronal function in a model of molecular pathology associated with schizophrenia. Mol Psychiatry 21, 1517–1526

105. Goversen, B., Jonsson, M. K., van den Heuvel, N. H., Rijken, R., Vos, M. A., van Veen, T. A., and de Boer, T. P. (2019) The influence of hERG1a and hERG1b isoforms on drug safety screening in iPSC-CMs. Prog Biophys Mol Biol

